# Heterogeneous genetic basis of age at maturity in salmonid fishes

**DOI:** 10.1101/2020.07.24.220111

**Authors:** Charles D. Waters, Anthony Clemento, Tutku Aykanat, John Carlos Garza, Kerry A. Naish, Shawn Narum, Craig R. Primmer

## Abstract

Understanding the genetic basis of repeated evolution of the same phenotype across taxa is a fundamental aim in evolutionary biology and has applications to conservation and management. However, the extent to which interspecific life-history trait polymorphisms share evolutionary pathways remains under-explored. We address this gap by studying the genetic basis of a key life-history trait, age at maturity, in four species of Pacific salmon (genus *Oncorhynchus*) that exhibit intra- and interspecific variation in this trait – Chinook Salmon, Coho Salmon, Sockeye Salmon, and Steelhead Trout. We tested for associations in all four species between age at maturity and two genome regions, *six6* and *vgll3*, that are strongly associated with the same trait in Atlantic Salmon (*Salmo salar*). We also conducted a genome-wide association analysis in Steelhead to assess whether additional regions were associated with this trait. We found the genetic basis of age at maturity to be heterogeneous across salmonid species. Significant associations between *six6* and age at maturity were observed in two of the four species, Sockeye and Steelhead, with the association in Steelhead being particularly strong in both sexes (*p* = 4.46×10^−9^ after adjusting for genomic inflation). However, no significant associations were detected between age at maturity and the *vgll3* genome region in any of the species, despite its strong association with the same trait in Atlantic Salmon. We discuss possible explanations for the heterogeneous nature of the genetic architecture of this key life-history trait, as well as the implications of our findings for conservation and management.

## Introduction

The repeated evolution of the same phenotype across taxa raises fundamental questions on how evolution might act on underlying genetic architecture, and how this architecture might constrain the phenotypic response (Elmer & Meyer, 2011). Well characterized examples of parallel evolution within individual species, such as adaptation to freshwater in stickleback fish, reveal the relative role of loci of varying effect sizes, as well as both ancestral and novel mutations (Jones et al., 2012; Liu, Ferchaud, Gronkjaer, Nygaard, & Hansen, 2018). For polymorphic traits that influence fitness, understanding how they are retained following species radiation addresses key questions on the predictability of evolution, particularly since speciation involves founder events that may act to reduce underlying genetic variation (Guerrero & Hahn, 2017). Three main but non-exclusive pathways for the retention of polymorphic traits during speciation events have been identified (Jamie & Meier, 2020); evolution on standing ancestral variation (Cortez et al., 2014), introgression (Giska et al., 2019), and novel mutations (Mundy, 2005). However, the extent to which interspecific life-history trait polymorphisms share evolutionary pathways remains relatively uncharacterized (Jamie & Meier, 2020). This knowledge can inform how evolution may act to maintain such variation within daughter species (Ayala & Campbell, 1974; Charlesworth, Nordborg, & Charlesworth, 1997; Guerrero & Hahn, 2017), how genetic architecture may predict the retention of such polymorphisms, and how related species may adapt to shared selection regimes (Jamie & Meier, 2020). Comparative analyses of the genetic basis of such traits between species also have implications for conservation, management, and food security. Shared genetic architectures that are well characterized in one taxon may inform the conservation of evolutionary processes in related species, and selective breeding programs (e.g. agriculture and aquaculture operations) in new species may take advantage of trait-linked markers previously identified in other species.

The age at which an individual reaches sexual maturity is a polymorphic life-history trait in many species that can have significant fitness consequences. For example, maturing later provides more opportunities to grow, potentially conferring larger body size and higher energy reserves at reproduction, which in turn can enable higher fecundity and increased offspring survival (Stearns, 1992). However, delayed maturity also increases the risk of mortality before reproduction and translates into longer generation times (Charnov & Gillooly, 2004; Kozlowski, 1992; Roff, Mostowy, & Fairbairn, 2002; Stearns, 1992). Given that this trait is generally heritable (Gjerde, 1984; Hankin, Nicholas, & Downey, 1993), the large variation in size and age at maturity observed within and among populations of many species indicates that there is often no single, optimal maturation strategy, and considerable research effort has focused on determining the ecological factors and evolutionary mechanisms leading to the maintenance of such variation across different species (Flatt & Heyland, 2011; Stearns, 1989). Further, in related species, there is some question about whether the same evolutionary processes result in shared trait variation. In addition to its fundamental importance to ecology and evolution, understanding the drivers of variation in size and age at maturity has applied relevance in fields such as fisheries (Conover & Munch, 2002), agriculture (Kenny, Heslin, & Byrne, 2018), aquaculture (Taranger et al., 2010), and conservation and management (Kindsvater, Mangel, Reynolds, & Dulvy, 2016). Yet, the genetic basis of variation underlying different maturation strategies across species has remained poorly understood until recently (Perry et al., 2014; Roff, 2011).

Fishes display more diversity in reproductive traits than any other vertebrate group (Mank & Avise, 2006). Within fishes, species in the family *Salmonidae* exhibit a wide range of this diversity (Schaffer, 2004), and the potential trade-offs between age at maturity (and therefore size) and survival are well established, particularly in anadromous species and populations (e.g. Fleming, 1996; Hankin et al., 1993; Healey, 1991). As growth rate is high during the marine migratory phase, variation in age may result in dramatic size differences, with the size of individuals potentially doubling with each additional year spent at sea. Larger, older individuals have been shown to have higher reproductive success once on the spawning grounds (Fleming, 1996; Janowitz-Koch et al., 2019; Mobley et al., 2019), but they also have a higher probability of mortality during the marine migration phase (Czorlich, Aykanat, Erkinaro, Orell, & Primmer, 2018), thus representing a classic evolutionary trade-off.

Variation in age at maturity is also highly relevant to the conservation and management of this socio-economically important family of fishes. First, variation in breeding age buffers populations from diversity loss following catastrophic environmental events via the ‘portfolio effect’ (Greene, Hall, Guilbault, & Quinn, 2010; Satterthwaite, Carlson, & Criss, 2017; Schindler et al., 2010). Second, such life-history variation reduces inter-annual variability in adult returns and the frequency of fishery closures, which benefits commercial, recreational, and subsistence fisheries and associated communities (Brown & Godduhn, 2015; Copeland, Ackerman, Wright, & Byrne, 2017; Greene et al., 2010; Schindler et al., 2010). Third, knowledge of age structure is important for modeling the effects of fisheries harvest and developing appropriate restoration strategies aimed at maintaining population diversity in fitness traits (Bowersox, Corsi, McCormick, Copeland, & Campbell, 2019; Hankin & Healey, 1986; Larsen et al., 2019; Ricker, 1980).

There is a pressing need to understand the mechanisms that drive variation in age at maturity in salmonids due to the trait’s ecological, evolutionary, and economic importance, particularly as the age structures of many populations have been changing in recent decades (Cline, Ohlberger, & Schindler, 2019; DeFilippo et al., 2019; Lewis, Grant, Brenner, & Hamazaki, 2015; Ohlberger, Ward, Schindler, & Lewis, 2018). Age at maturity has been shown to have high heritability in multiple species (Carlson & Seamons, 2008; Hankin et al., 1993; Tipping, 1991), suggesting significant genetic variation for this trait that may promote local adaptation (Vähä, Erkinaro, Niemela, & Primmer, 2008). The trait has also been shown to differ considerably between the sexes, with females maturing, on average, later than males (Fleming, 1996; Schaffer, 2004).

Recently, there have been significant advances in understanding the genetic basis of age at maturity in one salmonid species, Atlantic Salmon (*Salmo salar*), with a particular focus on the number of years spent in the marine migration phase (i.e. sea-age). Several studies have identified a large effect locus on chromosome Ssa25, with variation near a strong candidate gene, *vgll3*, explaining almost 40% of the variation in sea-age at maturity in wild-caught individuals from over 50 European populations, including several divergent lineages (Ayllon et al., 2015; Barson et al., 2015). Subsequent studies have also identified associations between the same genomic region and age for this species (Ayllon et al., 2019; Christensen, Gutierrez, Lubieniecki, & Davidson, 2017). Further, Barson et al. (2015) identified a second region on chromosome Ssa09 encompassing the candidate gene *six6* that was initially highly significant in a genome-wide association (GWA) analysis, although this signal was no longer statistically significant following correction for population structure. However, a recent study in an aquaculture population of Atlantic Salmon identified strong associations with early maturation and both the chromosome Ssa09 (*six6*) and chromosome Ssa25 (*vgll3*) regions (Sinclair-Waters et al., 2020), suggesting that the population stratification correction in Barson et al. (2015) may have been overly conservative. Interestingly, GWA studies of North American-origin Atlantic Salmon aquaculture stocks have failed to find strong associations between sea-age and variants in these regions (Boulding, Ang, Elliott, Powell, & Schaeffer, 2019; Mohamed et al., 2019), possibly due to a lack of variation in these regions (Sinclair-Waters et al., 2020). Given the associations identified in Atlantic Salmon and that both *six6* and *vgll3* have also been linked with pubertal timing in humans (e.g. Cousminer et al., 2013; Cousminer, Widen, & Palmert, 2016; Perry et al., 2014) and other mammals (Cánovas et al., 2014), these same loci may play a role in the maintenance of variation in age at maturity phenotypes in other salmonid species.

Pacific salmonids include numerous species in North America (genus *Oncorhynchus*), most of which exhibit anadromy (Behnke, 2002), and six of which show variation in the number of years spent in the marine environment prior to reproduction. Four species – Chinook Salmon (*O. tshawytscha*), Coho Salmon (*O. kisutch*), Sockeye Salmon (*O. nerka*), and Steelhead Trout (*O. mykiss*, the anadromous form of Rainbow Trout) – have been the focus of intense management and research. As a result, individual-level life-history data and tissue samples for genetic analyses are available for multiple populations in each of these species, making them a promising target for genotype-phenotype association studies. Chinook Salmon and Steelhead Trout exhibit the greatest variation in age at maturity (typically two to six years), Sockeye Salmon commonly mature at three to five years of age, and Coho Salmon between two and four years (Quinn, 2005), although populations can exhibit different age structures due to genetic (McKinney et al., 2019) and environmental factors such as local rearing and growing conditions (Hankin et al., 1993; Harstad, Larsen, & Beckman, 2014; Harstad et al., 2018; Taylor, 1990). Pacific salmon therefore provide an excellent opportunity to further assess the influence of *six6* and *vgll3* on age at maturity and improve our understanding of the extent to which genetic architectures are shared across species.

Here, we used a targeted single nucleotide polymorphism (SNP) approach to test for associations between the *six6* and *vgll3* genome regions and age at maturity in four Pacific salmonid species native to North America – Chinook Salmon, Coho Salmon, Sockeye Salmon, and Steelhead Trout - to determine whether these loci contribute to the maintenance of variation in age at maturity across related species. Samples were collected from two to six populations (hatchery and wild) of varying phylogenetic, phenotypic, and geographic backgrounds per species, and SNPs were identified in the *six6* and *vgll3* genes. Associations between these loci and age at maturity were then quantified using logistic and cumulative proportional odds models that accounted for potential confounding effects. For Steelhead Trout, we also tested for associations in other genomic regions with whole genome resequencing of groups with differing sea age, to determine whether additional regions were associated with this trait in this species. The results improve our understanding of the adaptive genetic variation underlying this key trait across multiple species and provide additional tools to monitor age structure in salmon populations of commercial, ecological, and evolutionary importance.

## Methods

### Sample collections

Populations of Chinook Salmon, Coho Salmon, Sockeye Salmon, and Steelhead Trout (hereafter referred to as Chinook, Coho, Sockeye, and Steelhead) for which individual-based collections of genetic samples and ages at maturity existed were identified for this study. Two to six populations, ranging from California to Alaska, were sampled per species and included those of hatchery and wild origin and variable age structures (Table 1, Tables S1-S4). Genomic DNA from most samples had been previously isolated. When needed, genomic DNA was extracted from tissue samples using DNeasy Blood & Tissue kits (Qiagen, Valencia, CA, USA) following the animal tissue protocol. Ages had been previously determined from a combination of coded wire and passive integrated transponder tags, growth rings on scales, and molecular pedigree analyses (Table 1). Ages of Chinook, Coho, and Sockeye correspond to age at maturity since these species are semelparous. Ages of Steelhead correspond to age at first reproduction since this species is iteroparous. There were five Steelhead samples from the Eel River population that matured at age six. Due to the low sample size, these individuals were combined with the age-five fish for association analyses. When possible, existing molecular pedigrees were utilized to ensure the sampling of unrelated individuals, as relatedness may bias results of association analyses (Korte & Farlow, 2013). Detailed population descriptions can be found in the Supplementary Information.

**Table 1.**
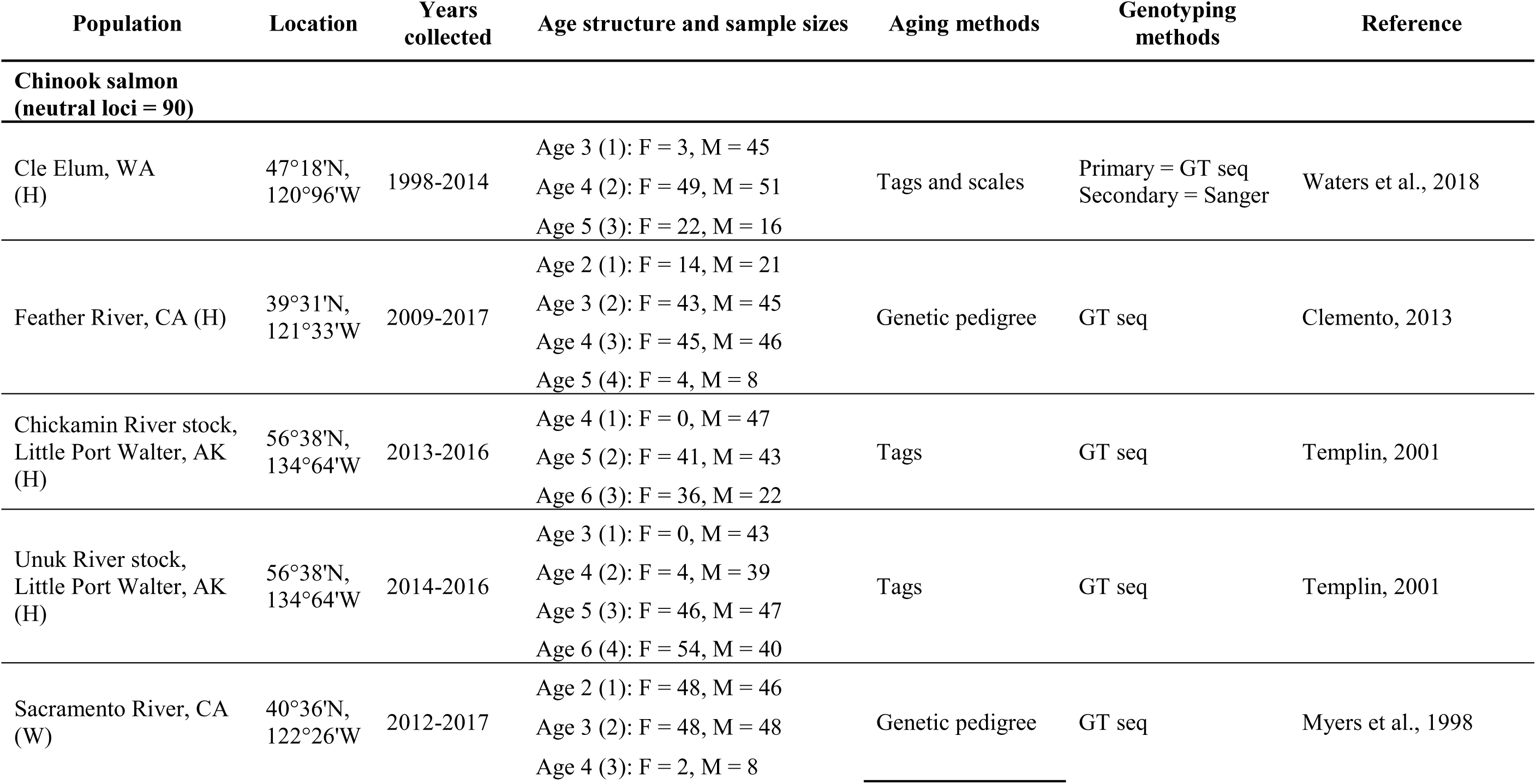
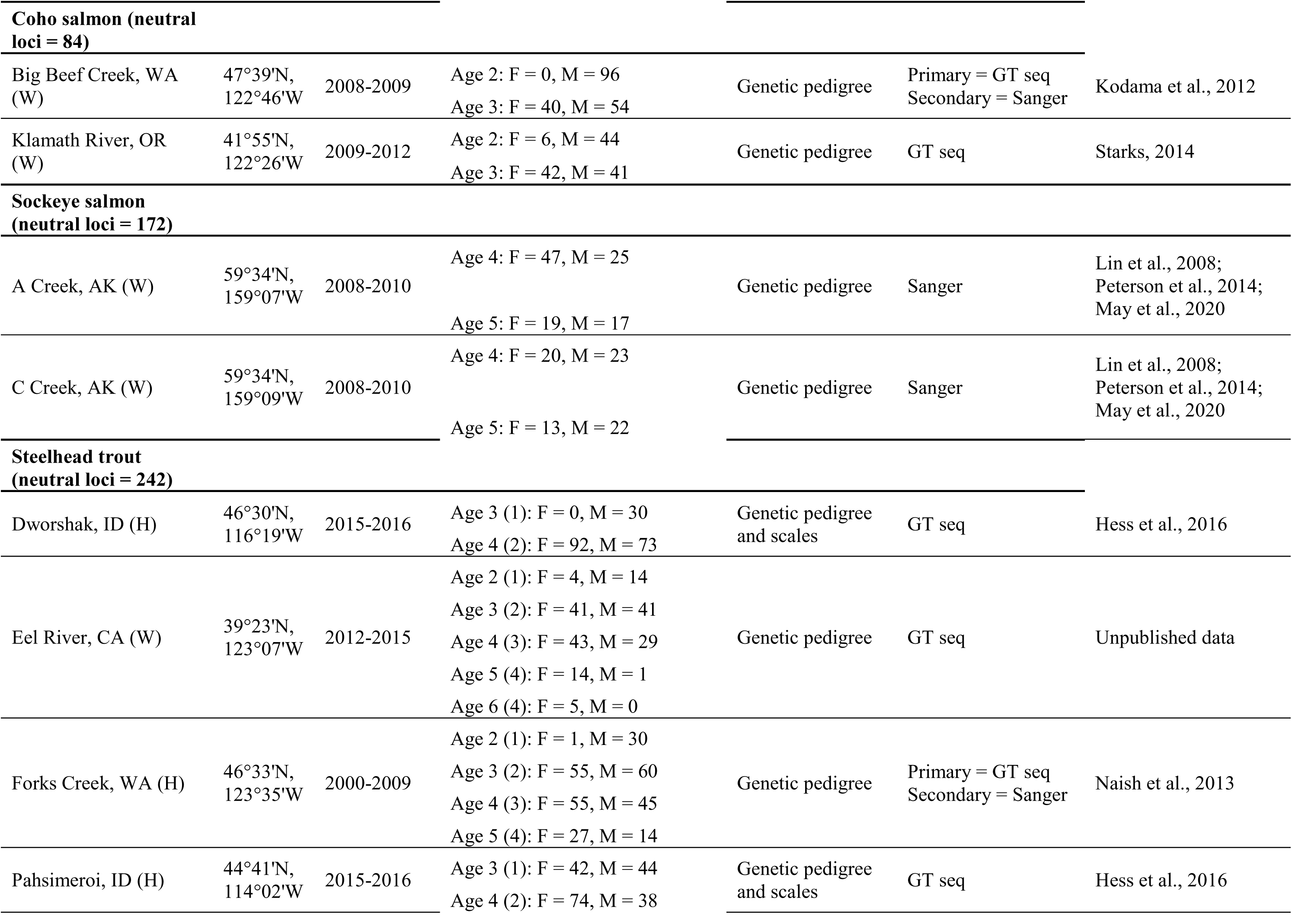
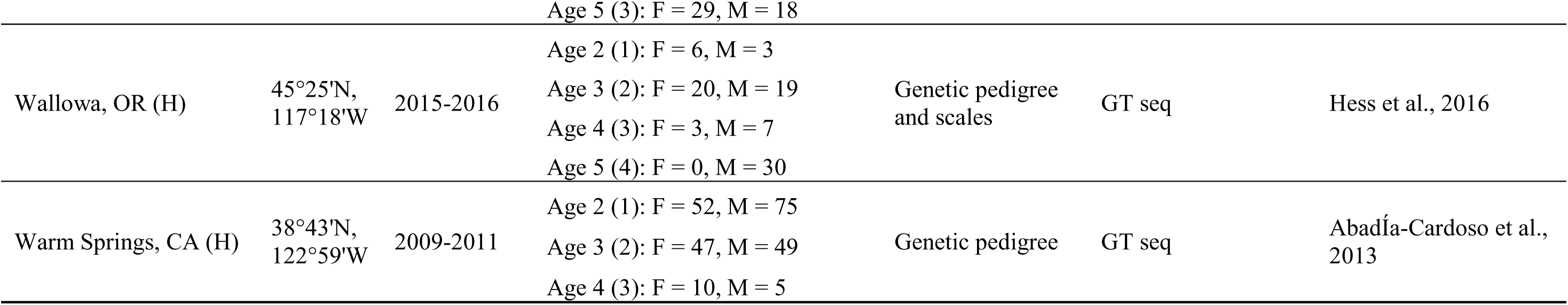
Populations of hatchery (H) and wild (W) Chinook Salmon, Coho Salmon, Sockeye Salmon, and Steelhead Trout that were analyzed in this study with sample sizes of females (F) and males (M) for each age class. Ages denote total age at maturity, with standardized ages (i.e. youngest age class of each population = 1) reported in parentheses for Chinook and Steelhead. Standardized ages were analyzed for these two species to account for inherent differences in age structure of the populations. The number of putatively neutral, polymorphic loci genotyped to serve as controls and assess potential confounding effects is also listed for each species. Population locations are given by State; CA - California, OR - Oregon, WA - Washington, AK - Alaska.

### Gene primer development

Sequences for the *six6* and *vgll3* genes in Atlantic Salmon were obtained from SalmoBase (v. 1, Samy et al., 2017) and aligned to the Coho Salmon (GenBank: GCA_002021735.1) and Rainbow Trout (GCF_002163495.1; Pearse et al., 2019) genomes using *Bowtie2* (v. 2.3.1, Langmead & Salzberg, 2012). The Chinook and Sockeye Salmon genomes were not explicitly considered during primer development, as they had not yet been released at the time of development. Local alignments, rather than end-to-end, were conducted in *Bowtie2* to account for potential differences in gene sequences between species, and up to ten alignments were reported (k=10) so that multiple alignments, including those to homeologous regions, could be investigated. Next, SAM output files from *Bowtie2* were converted to sorted, indexed BAM files using *samtools* (v. 1.3.1, Li et al., 2009). Alignments of the *six6* and *vgll3* genes to the Coho and Rainbow Trout genomes were then visualized using the Integrated Genome Browser tool (Freese, Norris, & Loraine, 2016). The preferred alignment of each gene was determined by the length and mapping quality (i.e. number of mismatches and indels) of the reported alignments, as well as *a priori* knowledge of chromosomal homologies between species obtained from comparative mapping (Kodama, Brieuc, Devlin, Hard, & Naish, 2014; Pearse et al., 2019).

Regions where a portion of the best alignments of the *six6* and *vgll3* Atlantic salmon gene sequences matched, or nearly matched, the Rainbow Trout and Coho Salmon genomes were candidates for primer development. For each gene, six candidate regions spanning approximately 580-780 base pairs (bp) were identified. Next, *Primer3* (Koressaar & Remm, 2007; Untergasser et al., 2012) was used to design PCR primers within 50-100 bp of the sequence tails; minimum, maximum, and optimal primer lengths were set to 21, 25, and 23 bp, respectively. The T7 (5’ TAATACGACTCACTATAGGG 3’) and T3 (5’ ATTAACCCTCACTAAAG 3’) universal promoter sequences were then added to all forward and reverse primers, respectively.

### PCR amplification

The six candidate regions for each gene were amplified using PCR in a subset of samples (*n*=8 per species) to evaluate amplification success, including possible targeting of gene duplications. 15 *u*L PCR reactions were performed, each comprising 1.5 *u*L of genomic DNA, 7.5 *u*L of 2X Qiagen Multiplex PCR Master Mix, 0.3 *u*L each of the newly-designed forward and reverse primers at 10 *u*M concentration, and 5.4 *u*L of pure water. The PCR protocol included: initial denaturation at 95°C for 15 minutes (min), ten cycles of [95°C for 30 seconds (s), 58°C for 90 s, 72°C for 90 s], 25 cycles of [95°C for 30 s, 62°C for 90 s, 72°C for 90 s], and a final extension at 72°C for five min. PCR products were cleaned using the ExoSAP-IT PCR Product Cleanup kit (Applied Biosystems) following the manufacturer’s protocol and then visualized using agarose gel electrophoresis. Primers that yielded single, high-quality bands of PCR product that matched the expected fragment lengths were deemed suitable for screening in a larger number of samples for single nucleotide polymorphism (SNP) identification.

### SNP identification and sample screening

Gene regions were amplified in 16 Chinook, 16 Coho, 16 Sockeye, and 16 Steelhead of varying age classes to identify SNPs. For this screening, 10 *u*L PCR reactions were performed, each comprising 1 *u*L of genomic DNA, 5 *u*L of 2X Qiagen Multiplex PCR Master Mix, 0.2 *u*L each of 10 *u*M forward and reverse primers, and 3.6 *u*L of pure water. The PCR protocol and PCR product cleaning were the same as in the initial amplification. Sanger sequencing (Macrogen, Inc.) was then used to sequence the cleaned PCR products.

*Geneious* (v. 10.1.3; https://www.geneious.com) was used to align sequences for each gene region and species using default parameters. SNPs were identified from the alignments, and emphasis was placed on SNPs with intermediate frequencies to maximize variability and the potential to detect genotype-phenotype associations. For Chinook, Coho, and Steelhead, consensus sequences were extracted for regions that contained SNPs; these sequences were then utilized for genotyping in two ways. First, a subset of populations - Cle Elum Chinook, Big Beef Creek Coho, and Forks Creek Steelhead - were genotyped using Sanger sequencing, and alignments were scored in *Geneious*. This initial data was then used to develop primers for high-throughput screening of all study samples using amplicon sequencing methods (Baetscher, Clemento, Ng, Anderson, & Garza, 2018; Campbell, Harmon, & Narum, 2015). All sequences of the *six6* and *vgll3* gene regions obtained from high-throughput screening were aligned and scored as in Baetscher et al. (2018). All Sockeye samples were genotyped directly from Sanger sequences rather than amplicons due to relatively small sample sizes.

SNPs were retained for final analyses if the minor allele frequency was greater than 10%, and individuals were included if they were genotyped successfully at one or more of the SNPs. When *six6* or *vgll3* contained multiple, highly correlated SNPs (based on Pearson’s r) that passed filtering, only one “target” SNP was tested. However, when possible, missing genotypes for the target SNP were imputed using the linked SNP(s). Steelhead samples from the Eel River, Forks Creek, and Warm Springs populations were the only ones genotyped for the *vgll3* SNPs, as there was not sufficient material remaining to genotype the Dworshak, Pahsimeroi, and Wallowa populations.

We also utilized 10 SNPs from the *six6* genome region in Steelhead that had been independently identified using a Pool-seq approach, described in full below (Schlötterer, Tobler, Kofler, & Nolte, 2014). Primers were developed with *Primer3* based on sequences surrounding each target SNP identified from the Pool-seq data. Primers were tested for amplification success in a multiplex reaction using the GT-seq protocol (Campbell et al., 2015). A total of 10 target SNPs were successfully developed (Table S13B) and produced reliable results for further genotyping following standard methods from Campbell et al. (2015). These 10 SNPs included six markers upstream of the *six6* gene on chromosome Omy25 that were presumably in regulatory regions from existing genome annotation, three markers in the *six6* gene sequence, and one SNP downstream of the *six6* gene.

Last, all samples were genotyped at suites of putatively neutral loci that had been previously developed for other research and monitoring purposes (Table 1; Tables S14-S17). The neutral loci served as “controls” during association testing and were also used to account for the possible confounding effects of population structure and relatedness in the association analyses.

### Association testing

Associations between age at maturity and SNPs within the *six6* and *vgll3* genome regions were tested using logistic regression and cumulative proportional odds models in R using the *stats* and *ordinal* packages, respectively (Christensen, 2019; R Core Team, 2019). Logistic regression models were employed for Coho and Sockeye since age at maturity was a binary trait for these species (Coho: ages two and three; Sockeye: ages four and five; Table 1). Cumulative proportional odds models were used for Chinook and Steelhead, as age at maturity in these species was an ordinal trait (Chinook and Steelhead: ages two to six; Table 1).

Age at maturity was modeled as a function of SNP, sex, and their interaction. To account for population stratification, a principal component analysis was conducted for each species using putatively neutral loci and the R package *adegenet* (v. 2.1.3, Jombart, 2008; Jombart & Ahmed, 2011). The PCs that reflected population structure (PC1 for Coho and Sockeye; PCs 1-4 for Chinook and Steelhead; Figures S1-S18), as well as their interactions with the SNP, were then included as covariates in the models. Relatedness between samples could not be explicitly included in the models. However, molecular pedigrees were utilized during study design to maximize the sampling of unrelated individuals for most populations (Table 1). Molecular estimates of pairwise relatedness between samples within each population were also generated to determine whether relatedness might have influenced the results (Supplementary Information).

The general form of each full model for Coho and Sockeye was: 

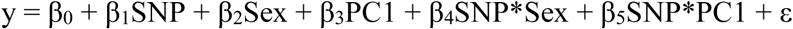

where y is the binary age at maturity (0 = younger age class, 1 = older age class), SNP is the genotype, Sex is the sex of the individual, and PC1 is the coordinate of the individual on principal component 1. Two full models were constructed for each species, one where the target SNP effect was modelled as additive (i.e. genotype treated as numeric, A/A=1, A/T=2, T/T=3) and a second where the SNP effect was non-additive (i.e. genotype treated as categorical instead of numeric). Models for Coho were evaluated using only males, as females were nearly invariant for age, and thus did not include sex as a covariate.

The general form of each full model for Chinook and Steelhead was: 

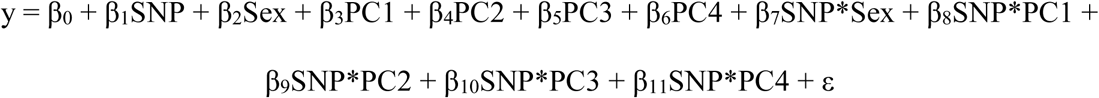

where y is the standardized age at maturity (i.e. youngest age class of each population = 1; Table 1; Table S5), SNP is the genotype, Sex is the sex of the individual, and PCs 1-4 are the coordinates of individuals along the first four principal components. Total age and a binomial age classification were also tested (Tables S5-S6). Four full models were constructed for these species, where the target SNP effect was modelled as additive or non-additive, and, within these models, the threshold structure was either constant or allowed to vary by sex since maturation thresholds may differ between the sexes. For all species, interactions between the SNP and other covariates were included in the full models to determine if SNP effects varied by Sex or location on the PCs (i.e. population). Additional interactions (e.g. Sex*PC1) were not included, as SNP effects were of primary interest.

Backward model selection was performed on all full models within a species (additive versus non-additive for all species; constant versus flexible threshold structure by sex for Chinook and Steelhead) using the *dredge* function of the *MumIn* package (Barton, 2016), and a preferred model was identified using corrected Akaike Information Criterion (AICc) scores. The significance of the SNP effect was then quantified by comparing the likelihood of the preferred model that included terms with the SNP to the likelihood of a model without the SNP effects. When AICc favored the non-additive SNP model, SNP effects were subsequently partitioned into additive and non-additive components and quantified using the same model comparison approach.

The significance of SNP effects may have been spuriously inflated by cryptic relatedness and residual population structure (Price et al., 2006) despite attempts to sample unrelated individuals and the incorporation of PCs into the models. To monitor and account for such genomic inflation, all polymorphic, putatively neutral loci (n=84 to 242 loci per species, Table 1) were evaluated using the same parsimonious model structures as the *six6* and *vgll3* SNPs. The significance of the *six6* and *vgll3* SNPs were then ranked among those obtained from the putatively neutral SNPs. Genomic inflation factors (i.e. lambda) were also estimated by dividing the median of the observed chi-square values by the median of the expected chi-square distribution of the same degrees of freedom (Devlin & Roeder, 1999; Marchini, Cardon, Phillips, & Donnelly, 2004). Chi-square test statistics of the *six6* and *vgll3* SNPs were then divided by the inflation factors, which were then used to obtain “corrected” *p*-values. It should be noted, however, that lambda values may be overly conservative due to the relatively few number of loci used to estimate them.

### Whole genome resequencing collections, library preparation, and sequencing

In order to assess whether other genome regions may harbor loci associated with age at maturity, whole genome resequencing was conducted in Steelhead. Specifically, whole genome resequencing of pooled sample collections was completed with six pools of returning adult male Steelhead that exhibited variable ocean age at maturation phenotypes and originated from three different locations (Table S9A). Individual fish for each pooled sample were collected at a fish trap at Bonneville Dam on the Columbia River (on the border between Oregon and Washington, U.S.A.), and the origin of their specific hatchery was identified through parentage analyses (see Hess et al., 2016 for details). Ocean age at maturation was determined by examination of scale annuli collected from each fish at Bonneville Dam, and total age was determined from parentage assignment results. Samples were grouped by ocean age at maturity phenotypes (one or two years in the ocean, or 1- and 2-ocean) within each population, with an average sample size of 48 male fish per pool (Table S9A). DNA was extracted individually with Chelex beads.

Phenotype pools within each population were prepared and sequenced using a standardized Pool-seq protocol described in Micheletti, Hess, Zendt, and Narum (2018). This pooled approach provides estimated allele frequencies for variants across the genome but not individual genotypes. Library preparation included normalizing individual DNA quantity using picogreen fluorescence on a Tecan M200 (Tecan Trading, AG, Switzerland). To ensure a similar contribution of each individual to a pool, individual DNA concentrations were not allowed to deviate more than 20% of the average DNA concentration of all individuals. For a given population, samples were fragmented with NEBNext Ultra dsDNA Fragmentase (New England Biolabs, Ipswich, MA, USA), pooled together, and filtered using Minelute purification (Qiagen, Venlo, Netherlands). Fragment end repair was performed with NEBNext Ultra End Prep (New England Biolabs, Ipswich, MA, USA), and fragments between 400-500 bp were selected for purification using a 25X AMPure beads solution (Beckman Coulter Inc, Indianapolis, IN). Fragments were then amplified with a NEBNext Ultra Q5 PCR protocol, cleaned with AMPure beads, and finally quantified with SYBR quantitative PCR (Thermofisher Scientific, Waltham, MA, USA). Amplified fragments were normalized and then sequenced with high-output runs on an Illumina NextSeq 500 (Illumina, San Diego, CA, USA) with paired-end 150bp reads (2×150bp).

### Whole genome resequencing bioinformatics and statistical analyses

For each Pool-seq library, raw 150 bp paired-end reads (2×150 bp) were processed using the PoolParty pipeline (Micheletti & Narum, 2018) that integrates several existing resources into a single pipeline. Briefly, this included multiple steps that started with trimming reads (to a minimum of 50 bp) with a quality score less than 20 using the trim-fastq.pl script part of Popoolation2 (Kofler, Pandey, & Schlotterer, 2011). Trimmed reads were then aligned to the *O. mykiss* genome assembly (GCF_002163495.1) using *bwa-mem* (Li, 2013) with default parameters. PCR duplicates were identified and removed using *SAMblaster* (Faust & Hall, 2014). SAMtools view module (Li, 2011) was used to sort BAM files, which were then combined using the SAMtools mpileup module that extracts SNP and coverage information for each pool. Best practices were implemented following Kofler et al. (2011) to filter out SNPs within 5 bp of indel regions from further analysis and those with minor allele frequency < 0.05. Variant positions were kept with a minimum of 15X depth of coverage and a maximum of 250X depth of coverage, which was intended to filter out regions that may be paralogs (and thus have high coverage) or regions that are likely represented by a small number of individuals (low coverage).

Filtered allele frequency data were then used to calculate fixation index (FST) between the collections comprising different ocean ages at maturity (1- or 2-ocean) within each population with a sliding window of 8000 bases with a step size of 100 bases. Genomic regions with statistically significant differentiation were determined using a local score technique that accounts for linkage disequilibrium of SNPs (Fariello et al., 2017). These local score analyses iteratively determined the tuning parameter (?) based on mean log10 *p*-values for each comparison, and significant regions were displayed (Bonferroni corrected α= 0.05) in the form of Manhattan plots using the R package *qqman* (Turner, 2014).

We also implemented a Cochran–Mantel–Haenszel (CMH) test that computes significance between groups of interest (Kofler et al., 2011); in this case, between pairs of 1-ocean and 2-ocean males from each of the three hatchery locations.

Genomic regions associated with differentiation in ocean age were deemed significant if they were shared between both local score analyses (analogous to a Bonferroni corrected α = 0.05) and the CMH test (Bonferroni corrected α = 0.05). These regions were then investigated for variant annotations using *SnpEff* (Cingolani et al., 2012) with *.gff files available for the genome assembly for Rainbow Trout (GCF_002163495.1).

## Results

### Gene primer development and PCR amplification

The preferred alignments of the Atlantic Salmon *six6* and *vgll3* gene sequences to the Coho Salmon and Rainbow Trout genomes were located on the chromosomes predicted by comparative mapping (Tables S10, S12). Gene sequences also aligned to other chromosomes, including homeologous chromosomes (e.g. Co26 in Coho and Omy22 in Rainbow Trout for *vgll3*), but the alignments were shorter and had lower quality (i.e. more mismatches and indels; Table S10). Six candidate regions for PCR amplification and SNP discovery were identified throughout each gene based on these alignments (Table S11A-B), and most regions amplified successfully.

### SNP identification and sample screening

Three regions from *six6* and *vgll3* were then screened in a larger number of samples (n=16 of varying ages per species) to identify SNPs and to generate consensus sequences (Table S11C-D) for further SNP development using amplicon sequencing methods. After genotyping and filtering for minor allele frequency, one to two target SNPs were identified for association analyses within each gene for Chinook, Coho, and Sockeye (Tables S1-S4, S13A). Five SNPs were identified within the *vgll3* gene for Steelhead, and there were 10 SNPs within the *six6* gene identified from the Pool-seq approach (Table S13B). For species-gene combinations with multiple target SNPs, we only present results for the SNP with the strongest association; results for other SNPs are in Table S6. Samples were also genotyped at putatively neutral loci (n=84 to 242 loci per species, Table 1; Tables S1-S4, S14-S17).

### Association testing in species with two age classes

Age at maturity in male Coho Salmon was not significantly associated with *six6* or *vgll3* (Table 2; Figure 1; Table S7). In contrast, *six6* was significantly associated with age in Sockeye Salmon, before and after correcting for potential genomic inflation (*p*=0.008, Table 2; Figure 1; Table S7). When partitioned, the non-additive SNP effect was significant while the additive effect was marginally non-significant (Table 2). The probability of delaying maturation (i.e. mature at age five rather than at age four) was 0.34 and 0.53 for the two alternative homozygous genotypes in female Sockeye, while those for male Sockeye were 0.48 and 0.66, respectively (Figure 1). That is, female and male Sockeye Salmon with the C/C genotype were 1.56 and 1.38 times more likely to mature at age five than individuals with the A/A genotype (Figure 1; Table S8A). There was no significant difference in maturation probability between the two sexes. The association of *six6* with age at maturity in Sockeye was greater than any of the 172 putatively neutral SNPs (Table 2).

**Table 2.** Results of association analyses between age at maturity and the *six6* and *vgll3* genes for each species based on the most parsimonious models. Multiple SNPs within the *six6* and *vgll3* gene regions were tested for Chinook and steelhead. Results reported here are for the SNPs with the most highly-associated additive components; those for the other SNPs are reported in Table S6. SNP rank indicates the relative significance of the *six6* and *vgll3* SNP associations compared to all putatively neutral SNPs that were tested in the same models. Significant *p*-values are in bold.

| Gene | Species | N, {Age classes}<sup>1</sup>, N<sub>pop</sub> | Parsimonious model<sup>2</sup> | AICc with SNP | Overall (adjusted<sup>3</sup>) SNP *p*-values | Lambda | Additive (non-additive<sup>4</sup>) component *p*-values | SNP Rank | Model | Notes |
| --- | --- | --- | --- | --- | --- | --- | --- | --- | --- | --- |
| *six6* | Chinook | 1042, {1-4}, 5 | *six6* <sub>allelic</sub> + *PC1* + *PC2* + *PC3* + *PC4* + *six6* <sub>allelic</sub> * *PC4* | 2453.2 | **0.002** (0.075) | 2.46 | N/A | 11 / 91 | Ordinal, nominal = sex | one of two SNPs tested |
|  | Coho | 233, {2-3}, 2 | *six6*<sub>allelic</sub>+*PC1* | 313.1 | 0.104 (0.319) | 2.65 | N/A | 21 / 85 | Binomial | Only males<sup>5</sup> |
|  | Sockeye | 176, {4-5}, 2 | *six6*<sub>geno</sub> + *Sex* | 226.9 | **0.003 (0.008)** | 1.22 | 0.093 (**0.003**) | 1 / 173 | Binomial |  |
|  | Steelhead | 1207,{1-4}, 6 | *six6* <sub>geno</sub> + *PC1* + *PC2* + *PC3* + *PC4* + *Sex* + *six6* <sub>geno</sub> * *PC1* + *six6* <sub>geno</sub> * *PC2* + *six6* <sub>geno</sub> * *Sex* | 2633.4 | **1.835E-19 (4.456E-9)** | 1.94 | **2.664E-16 (0.002)** | 1 / 243 | Ordinal, nominal = sex | SNP Omy25_61294400 |
| *vgll3* | Chinook | 1046, {1-4}, 5 | *vgll3* <sub>geno</sub> + *PC1* + *PC2* + *PC3* + *PC4* + *Sex* + *vgll3* <sub>geno</sub> * *PC2* + *vgll3* <sub>geno</sub> * *PC3* + *vgll3* <sub>geno</sub> * *PC4* + *vgll3* <sub>geno</sub> * *Sex* | 2460.7 | **8.286E-6** (0.147) | 2.86 | **0.028 (3.425E-5)** | 23 / 91 | Ordinal, nominal = sex | one of two SNPs tested |
|  | Coho | 203, {2-3}, 2 | *vgll3*<sub>allelic</sub> + *PC1* | 274.9 | 0.856 (0.900) | 2.11 | N/A | 83 / 85 | Binomial | Only males<sup>5</sup> |
|  | Sockeye | 160, {4-5}, 2 | *vgll3*<sub>geno</sub> | 203.2 | **0.047** (0.095) | 1.30 | 0.804 (0.071) | 12 / 173 | Binomial |  |
|  | Steelhead | 656,{1-4}, 3 | *vgll3* <sub>allelic</sub> + *PC1* + *PC2* + *PC4* + *Sex* + *vgll3* <sub>allelic</sub> * *PC1* + *vgll3* <sub>allelic</sub> * *PC2* | 1483.4 | **0.016** (0.185) | 2.14 | N/A | 48 / 243 | Ordinal | SNP *vgll3*.1_309 |

<sup>1</sup>age classes for coho and sockeye salmon represent total age at maturity, while those for Chinook and steelhead have been standardized (i.e. youngest age class of each population = 1) to account for inherent differences in age structure between populations.
<sup>2</sup>geno and allelic refers to non-additive and additive modelling of genotypes, respectively. Null models do not include any terms with the SNP.
<sup>3</sup>*p*-values after correcting for genomic inflation (i.e. lambda).
<sup>4</sup>additive vs. non-additive effects were partitioned when a non-additive model was preferred by model selection; these $p$ -values are unadjusted for possible genomic inflation.
<sup>5</sup>age at maturity was nearly fixed in females: six age 2 fish and 82 age 3 fish

**Figure 1.**
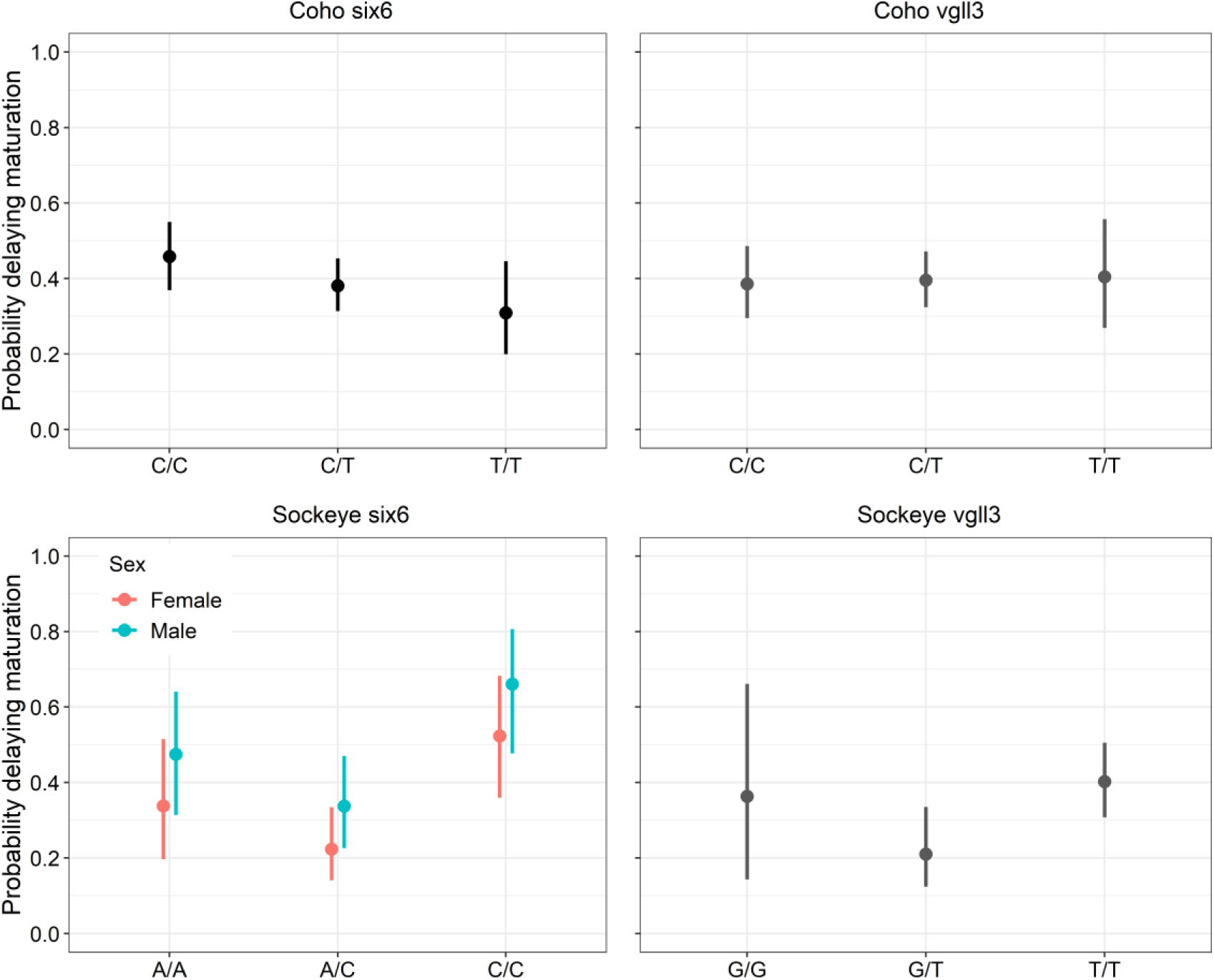
Predicted probabilities of delaying maturation with 95% confidence intervals for male Coho Salmon and Sockeye Salmon of both sexes for the genotypes of target SNPs in the *six6* and *vgll3* genome regions, obtained from the logistic regression models. When the most parsimonious model included PCs, the mean PC value was used when calculating probabilities. Only male Coho Salmon were analyzed, as females were nearly invariant in age. Probabilities for Sockeye Salmon at *vgll3* apply to both sexes since sex was not included in the most parsimonious model.

*Vgll3* was also significantly associated with age at maturity in Sockeye Salmon, although the effect became non-significant after accounting for genomic inflation (Table 2; Table S7). Further, the probabilities of delaying maturation were not significantly different between the two homozygous genotypes (Figure 1; Table S8A). Eleven neutral SNPs exhibited stronger associations with age than *vgll3* in Sockeye Salmon.

### Association testing in species with four age classes

Both *six6* and *vgll3* were significantly associated with standardized age at maturity in Chinook Salmon (Table 2; Figure 2; Table S7). However, the SNP effects became non-significant after accounting for potential genomic inflation. The cumulative probabilities of maturation were not significantly different between the two homozygous genotypes in both gene regions, although the maturation probabilities of male Chinook with the T/T genotype appeared to be lower, albeit non-significant, for the younger age classes than the G/G genotype in *vgll3* (Figure 2; Table S8B). In addition, the *p*-values for 10 and 22 putatively neutral loci were smaller (i.e. more significant) than the *six6* and *vgll3* genes, respectively. The specific effects of *vgll3* in Chinook Salmon may vary by population, as suggested by the interactions of the SNP with PCs 2-4. The influence of *vgll3* decreased as the location of individuals along PC2, PC3, and PC4 increased (Table S7). However, there is no apparent geographic gradient in population separation along PCs 2-4 (Figures S2-S5), so extrapolating these effects to other populations may be difficult. The overall effects of *six6* and *vgll3* did not differ significantly between the sexes of Chinook Salmon.

**Figure 2.**
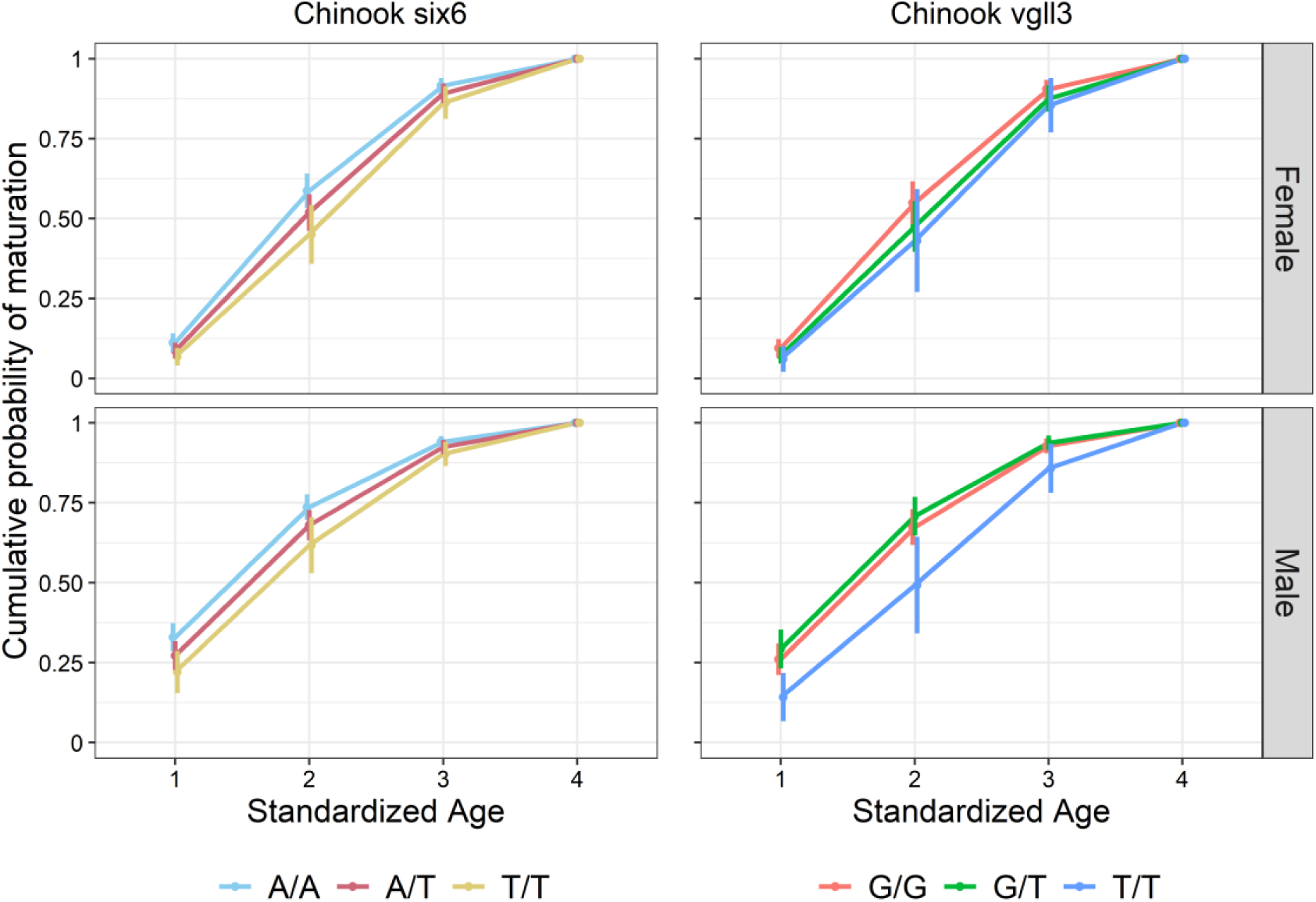
Cumulative probabilities of maturation with 95% confidence intervals for each standardized age class of male and female Chinook Salmon for the genotypes of target SNPs in the *six6* and *vgll3* gene regions, obtained from a cumulative proportional odds model. Mean PC values were used when calculating probabilities. As standardized age four was the oldest age, cumulative probabilities of maturation equal one for this class (i.e. all fish must mature by standardized age four).

*Six6* was highly associated with age at maturity in Steelhead Trout (*p* = 4.46×10^−9^ after adjusting for genomic inflation), with additive effects being greater than non-additive effects (Table 2). The probabilities of maturing at younger ages were higher for the G/G genotype than the A/A genotype (Figure 3). Specifically, female Steelhead with the G/G genotype were 2.53 and 1.34 times more likely to mature by age one and by age two, respectively, than fish with A/A genotype; male Steelhead with G/G were 2.96 and 1.34 times more likely to mature by age one and by age two than those with A/A (Figure 3; Table S8B). While the maturation thresholds varied by sex, the effect of *six6* on age did not differ significantly between females and males (Table S7). There was a significant interaction between *six6* and PC2, which separated the Forks Creek population from all others (Figure S17); the probability of delayed maturation increased slightly as the position along PC2 increased (Table S7). The other nine SNPs within *six6* were also significantly associated with age at maturity (Table S6), and all 10 *six6* SNPs exhibited stronger associations with age at maturity than the 242 putatively neutral SNPs tested (Table 2).

**Figure 3.**
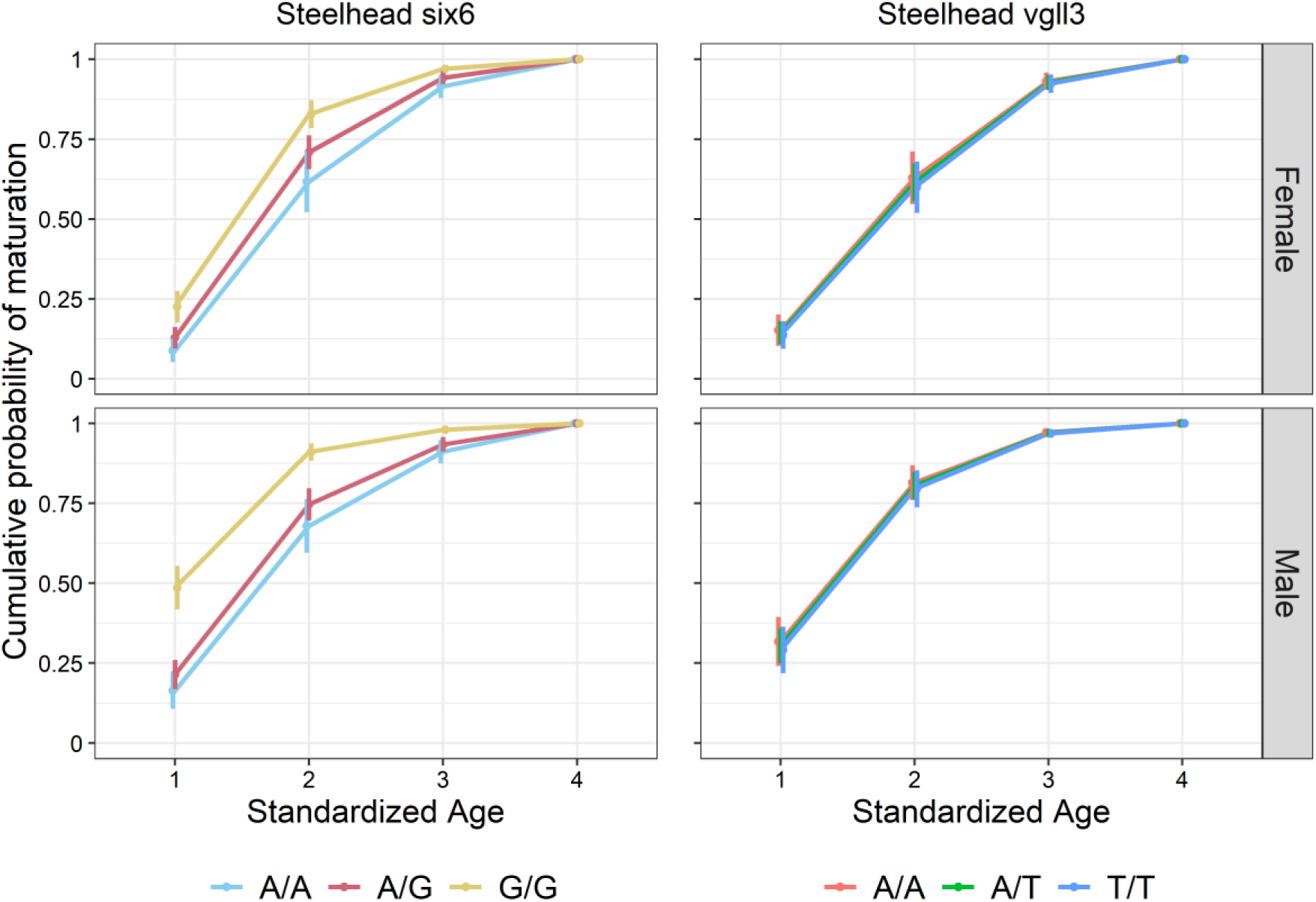
Cumulative probabilities of maturation with 95% confidence intervals for each standardized age class of male and female Steelhead Trout for the genotypes of target SNPs in the *six6* and *vgll3* gene regions, obtained from a cumulative proportional odds model. Mean PC values were used when calculating probabilities. As standardized age four was the oldest age, cumulative probabilities of maturation equal one for this class (i.e. all fish must mature by standardized age four).

Similar to Sockeye Salmon, *vgll3* was significantly associated with age in Steelhead Trout, but the effect became non-significant after correcting for genomic inflation (Table 2). The cumulative maturation probabilities of the three *vgll3* SNP genotypes were very similar (Figure 3; Table S8B), indicating that age was not strongly correlated with variation at *vgll3*. Last, 47 of the 242 neutral SNPs exhibited stronger associations with age than *vgll3* (Table 2).

### Whole genome resequencing in Steelhead Trout

Whole genome resequencing provided a means of assessing whether other genome regions may harbor loci associated with age at maturity. After filtering, the six male libraries representing paired collections (one versus two years in the ocean) from three populations had sequence coverage greater than 15X for a range of 48.6-67.0% of the genome (mean 62.2%; Table S9A). The mean depth of coverage for the six libraries was 23.0X with a range of 20.2-25.6X. There were a total of 2,850,964 SNPs identified across all six libraries, and allele frequencies were estimated for each variant in each of the six collections.

Tests for differences in allele frequencies of each paired collection of males (one versus two years in the ocean) within each of the three populations revealed several SNP markers on chromosome Omy25 that were significantly different for each pair based on FST and Fisher’s exact tests. Additionally, CMH tests that compared pools of different ages at maturity for each of the three population pairs identified a region on chromosome Omy25 as consistently significant (Table S9B). Finally, local score results that account for multiple SNPs in physical linkage demonstrated that the signal on chromosome Omy25 remained as the strongest signal (*p* < 0.001 after Bonferroni correction; Figure 4; Table S9B). However, two regions on chromosomes Omy15 and Omy26 were also identified from local score results but with much lower levels of significance (Table S9B). The highly significant region on chromosome Omy25 included markers that were located within and upstream of *six6*, with a total of 216 SNPs between positions 61.27-61.32Mb (Figure 4).

**Figure 4.**
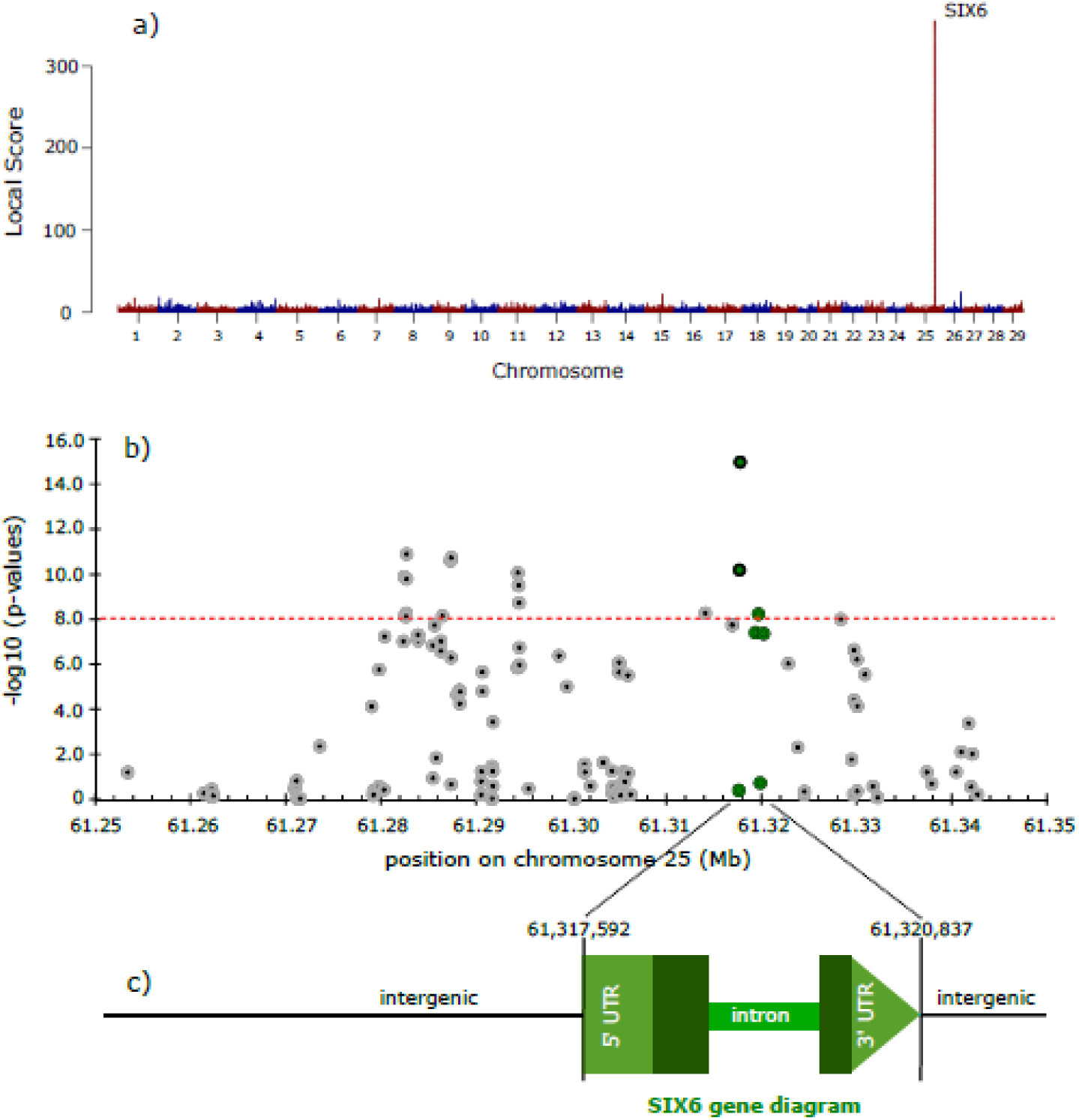
Genome-wide association test for Steelhead Trout including a) Manhattan plot of genomic regions associated with ocean-age at maturity in paired collections (1-ocean vs. 2-ocean) of male Steelhead from three populations (Local score results from CMH test of each pair of collections per location); b) Manhattan plot (*p*-values from Fishers exact test) zoomed to the significant region on chromosome Omy25 with green SNPs from the *six6* gene and gray SNPs intergenic, dashed red line is Bonferroni-corrected significance threshold; c) gene diagram for *six6*.

## Discussion

We examined whether two genes known to influence age at maturity in Atlantic Salmon (genus *Salmo*), *six6* and *vgll3*, also contribute to the phenotypic variation in this trait within four species of Pacific salmonid fishes (genus *Oncorhynchus*). We also examined the broader genetic architecture of this trait in one species, Steelhead Trout, to determine the relative genetic contribution of these genes to phenotypic variation compared to other loci. Two to six populations representing different phylogenetic, phenotypic, and geographic backgrounds were sampled per species. We found significant associations between *six6* and age at maturity in both Sockeye Salmon and Steelhead Trout after correcting for genomic inflation, with the association in Steelhead being particularly strong. Indeed, the odds of maturing by certain ages in this species differed by factors of 1.34 to 2.96 between the two alternative homozygotes, and those in Sockeye differed by factors of 1.38 to 1.56. In contrast, while *vgll3* has been found to be strongly associated with age at maturity in Atlantic Salmon, no significant associations were detected in the four Pacific salmonid species following correction for genomic inflation. Our examination of the relative contribution of these loci in one species, Steelhead, using whole genome resequencing also revealed the role of additional loci on separate chromosomes, although the significance of the associations were lower than *six6*. We sought to expand our understanding of the extent to which the genetic architecture of ecologically and evolutionarily important polymorphic traits may be conserved across species radiations. Combined, our results indicate that there is a heterogeneous genetic basis for age at maturity in salmonid fishes, including shared evolutionary pathways for age at maturity involving polymorphisms in the *six6* genomic region in at least two Pacific salmon species, but no clear evidence for *vgll3* being linked with this trait in the genus *Oncorhynchus.*

The presence of significant associations in both Pacific and Atlantic salmonids suggests that the influence of *six6* on age at maturity has been at least partially conserved over 21 million years of evolution (Lien et al., 2016). Given that significant associations were observed between *six6* and age at maturity in two populations of Sockeye and six populations of Steelhead, this study provides evidence that this genome region harbors broad, ecologically significant variation in salmonid fishes. The *six6* region has frequently been identified as an outlier in numerous evolutionary genetic studies in Atlantic Salmon (summarized in Prichard et al. 2018) and has been associated with landscape factors related to flow rate (Pritchard et al., 2018) as well as the run-timing life-history trait (Cauwelier, Gilbey, Sampayo, Stradmeyer, & Middlemas, 2018). Further, a RAD locus associated with spawning site selection in Sockeye Salmon (Veale & Russello, 2017) was identified to be located very close to the *six6* genome region (Pritchard et al., 2018). However, associations between *six6* and age at maturity have been inconclusive until recently. Early studies in Atlantic Salmon suggested that strong associations were inflated, or even entirely driven, by population structure (Barson et al., 2015; Johnston et al., 2014). Yet, a recent study detected a strong association between *six6* and early maturation (one year post smoltification, also known as ‘grilsing’) in >11,000 male Atlantic salmon from a single aquaculture strain, where population stratification is unlikely to be an issue (Sinclair-Waters et al., 2020). The results reported here strengthen the evidence that this locus is relevant to polymorphisms in age at maturity in additional salmonid species.

The molecular functions of *six6* uncovered thus far in vertebrates provide insight into the range of potential functional mechanisms that could link genotype and phenotype in salmonids. The sine oculis homeobox (SIX) protein family consists of evolutionarily conserved transcription factors found in organisms from flies to vertebrates (reviewed by Kumar, 2009). Specific functions of *six6* include playing a role in eye development (Seo, Drivenes, Ellingsen, & Fjose, 1998) and circadian timing in both mammals and fish (Clark et al., 2013; Watanabe et al., 2012). *Six6* has also been identified as a key fertility regulator in mammals (Larder, Clark, Miller, & Mellon, 2011) and has been linked with human pubertal timing (Hou et al., 2017; Perry et al., 2014). Further, *six6* has reported roles in the establishment of the brain-pituitary-gonad (BPG) axis in vertebrates (Jean, Bernier, & Gruss, 1999; Seo et al., 1998). Gene expression studies in Atlantic salmon support the known roles of *six6* in BPG axis signaling in the hypothalamus and pituitary, as well as eye development, and also suggest links with the Hippo signaling pathway (Kurko et al., 2020). The Hippo pathway is known to regulate cell fate commitment and organ (including gonadal) growth (Kjaerner-Semb et al., 2018; Li et al., 2015). Interestingly, *vgll3* is also known to be involved in the Hippo pathway (Kjaerner-Semb et al., 2018; Kurko et al., 2020), suggesting that genes in this pathway may be relevant to several ecologically important traits in salmonid fishes. However, linking genes of this pathway more directly with the ecological traits of interest awaits further study.

The *vgll3* gene has been linked to age at maturity in Atlantic Salmon in a number of independent studies (Ayllon et al., 2015; Ayllon et al., 2019; Barson et al., 2015; Debes et al., 2019; Verta et al., 2019), as well as with pubertal timing in humans and other mammals (e.g. Cánovas et al., 2014; Cousminer et al., 2013; Cousminer et al., 2016; Perry et al., 2014). The conserved nature of this gene region-phenotype association led us to hypothesize that the same region may be associated with age at maturity in Pacific salmonids. However, our results did not provide conclusive evidence to support this hypothesis. Although *vgll3* genotypes were non-randomly associated with age at maturity in Chinook, Sockeye, and Steelhead, the associations became non-significant after correcting for genomic inflation. Additionally, in each species, a number of randomly chosen SNPs across the genome exhibited even stronger associations with age, suggesting that any effect of *vgll3* is, at best, minor in the Pacific salmon populations assessed in this study.

There are several possible biological explanations for the heterogeneous results observed between the four Pacific salmon species analyzed here, as well as between Pacific and Atlantic salmon. First, the age at maturity trait may have evolved a different genetic architecture in some or all of the species investigated, a point discussed in greater detail below. Further, while salmonid fishes are known to exhibit local adaptation (Fraser, Weir, Bernatchez, Hansen, & Taylor, 2011; Taylor, 1991), including for age at maturity (Quinn, Wetzel, Bishop, Overberg, & Rogers, 2001; Roni & Quinn, 1995), there is also substantial variability in the degree of local adaptation in different traits among populations (Fraser et al., 2011), and age at maturity may not always be a key trait involved.

Differences in the degree to which age at maturity is under sexual conflict and how it is resolved may have also contributed to the heterogeneous results across species. Age at maturity is under sexual conflict in many salmonid species, whereby the optimal maturation age differs for males and females. One common evolutionary solution to resolve sexual conflict in other taxa is for genes controlling traits subjected to sexual conflict to reside on the sex chromosomes, which more readily allows for sex-specific expression patterns to evolve (Mank, 2017). Although they have a sex-determining locus (*sdY*; Yano et al., 2013), salmon do not have morphologically differentiated sex chromosomes, and therefore other evolutionary solutions are needed. In Atlantic Salmon, the sex-dependent dominance observed at the *vgll3* locus has been suggested to contribute to the partial resolution of sexual conflict (Barson et al., 2015). In Chinook salmon, it was recently reported that age at maturity in males is associated with male-specific haplotypes from a region on Ots17 (McKinney, Nichols, & Ford, 2020; McKinney et al., 2019). While this region may include the sex determining locus *sdY*, McKinney et al. (2020) also showed that the sex determination region can be translocated between two chromosomes, Ots17 and Ots18. Age at maturity loci have also been linked to the sex chromosome in Coho Salmon, the sister species to Chinook Salmon, in QTL mapping studies (Kodama, Hard, & Naish, 2018). Linkage of a large effect gene or suite of haplotypes controlling age at maturity with the sex determining region may be an effective strategy for intra-locus sexual conflict resolution, but the specific gene(s) involved are still unknown. Further, it remains to be determined whether similar mechanisms may occur in additional populations and species of salmonids.

Third, only one of the four species studied here, Steelhead Trout, exhibits similar age at maturity characteristics compared to Atlantic Salmon, as both species are iteroparous and exhibit more than two maturation ages. Interestingly, Steelhead Trout was also the species where the strongest single locus association with age at maturity was observed, albeit at *six6*. Yet, *six6* was also significant in Sockeye Salmon, a species not known to be iteroparous (Quinn, 2005), and this association was detected in two populations compared to the larger number of populations surveyed here for Steelhead. The age structure in Coho Salmon was the simplest within species studied here – only the males were polymorphic for this trait, and there were only two age classes. However, a broader range of species and populations needs to be studied before it can be concluded whether or not certain life-history characteristics (e.g. iteroparity) may be important in determining the genetic architecture of age at maturity.

There may also be technical explanations for the heterogeneous results we observed that are associated with study design. Specifically, we relied on populations with extensive, individual-based phenotypic and genetic collections. As such, Coho and Sockeye Salmon were represented by only two populations each, while Chinook Salmon and Steelhead Trout represented very diverse lineages. The power to detect associations may have been reduced in species with few populations, although we note that a significant association with *six6* was detected in Sockeye Salmon. Lineage divergence in interior and coastal populations of Chinook Salmon and Steelhead Trout is substantial (Nielsen, 1999; Waples, Teel, Myers, & Marshall, 2004), reflecting past glaciation events that may have resulted in the evolution of different architectures underlying the trait. Efforts to reduce the influence of population structure were implemented here; nevertheless, trait-associated loci are known to have population-specific effects in Chinook Salmon (McKinney et al., 2020). In fact, our results also suggest population-specific effects. Our study also included both hatchery and wild fish. Hatcheries have been known to truncate age at maturity (Larsen et al., 2019), and different selection regimes and environments of hatcheries might influence the genetic architecture underlying the phenotype (McKinney et al., 2020). We addressed the former by standardizing age at maturity between populations, but acknowledge that the latter may have played a role in our study.

Differences in results between species might also be associated with the genotyping. Specifically, surveys were not conducted over the entire genome. Instead, we used a relatively small number of putatively neutral SNPs for estimating genome inflation and correcting for population structure, which may have influenced the statistical power of the analyses. Similarly, relatedness could not be explicitly incorporated into the models. However, we took advantage of known pedigrees where possible to select unrelated individuals, and molecular estimates of pairwise relatedness suggest that fish within populations were predominately unrelated. The fact that a full genome-wide analysis was conducted in only one of the four species means that additional loci that influence age at maturity were not taken into account in three species, although it appears that *six6* has had a significant role in explaining the divergence between phenotypes in Steelhead compared to the other two loci detected on chromosomes Omy15 and Omy26.

Despite these caveats, the results presented here provide an intriguing insight into the evolution of a key trait that is polymorphic across Atlantic and Pacific salmonids. We can conclude that while the genetic basis of age at maturity appears to be heterogeneous both between and within salmonid genera, phenotypic polymorphisms associated with *six6* represent shared evolutionary processes across the two genera, although at this stage we cannot identify the exact mechanisms involved (Jamie & Meier, 2020). It is interesting to examine the relationships within Pacific salmonids within this context (Crete-Lafreniere, Weir, & Bernatchez, 2012; Macqueen & Johnston, 2014). Steelhead is the most divergent (i.e. outgroup) of the species studied here. This means that the common ancestor for Steelhead Trout and Sockeye Salmon, the two species that share the trait association with *six6*, pre-dates the divergence of all four species, and so the role of the locus has persisted throughout their radiation. If *six6* is indeed not associated with age at maturity in the remaining species examined, then it implies that alternative pathways have evolved. The evolution of quantitative traits depends on the relative role of large and small effect loci, their interactions, and the nature of selection acting upon them (Barton & Keightley, 2002; Kardos & Luikart, 2020; Oomen, Kuparinen, & Hutchings, 2020). For example, simulation studies have suggested that the evolution of large effect genes may only be favored under conditions where there is strong selection toward different (local) optima combined with moderate levels of gene flow (Yeaman & Whitlock, 2011). Rates of straying (i.e. gene flow) in salmonids vary between species, populations, and even life-history types, with those of Coho and Sockeye Salmon being, on average, lower than Atlantic Salmon, Chinook Salmon, and Steelhead Trout (Keefer & Caudill, 2014), and levels of gene flow may have played a role in the evolution of the trait within individual species. It has also been shown that the molecular basis of ecologically relevant traits is heterogeneous and does not always involve conserved mechanisms. For example, different genes in the same molecular pathways have been linked with similar phenotypes (Kronforst et al., 2012). As a transcription co-factor, *vgll3* expression likely influences a number of other genes (Kurko et al., 2020; Simon, Faucheux, Zider, Theze, & Thiebaud, 2016), and thus it is possible that variation in a different gene or genes in the same pathway could be more directly linked with age at maturity in some of the species investigated here. Indeed, *six6* may also be such a gene.

The discovery of genetic variation underlying ecologically important phenotypic traits across species has significant potential to inform their conservation and management, because architectures that predict evolution in one might be applied to the other. Age at maturity has been declining in many species of Pacific salmonids over recent decades (Bowersox et al., 2019; Cline et al., 2019; Lewis et al., 2015; Ohlberger et al., 2018), leading to lower reproductive potential of females (Ohlberger et al., 2020) and decreased population resilience through reduced phenotypic diversity (Greene et al., 2010; Schindler et al., 2010). Although *six6* is a candidate gene significantly associated with age at maturity in Sockeye and Steelhead, markers developed from this study need to be validated more broadly to determine population and sex-specific effects across the geographic distribution of each species, and the relative contributions of other loci to phenotypic variance need to be identified. In populations where markers from *six6* effectively predict phenotypic variation for age at maturity, it is possible to use this information to better understand mechanisms underlying observed phenotypic changes collected across temporal and spatial replicates, including monitoring anthropogenic effects such as fisheries-induced selection and supportive breeding. Monitoring of both adaptive and neutral (e.g. Schwartz, Luikart, & Waples, 2007) genetic variation might inform management actions such as harvest regulations aimed at conserving a broad portfolio of life-history variation that can be vital for species to persist in dynamic ecosystems (e.g. Schindler et al., 2010). While acknowledging that gene-targeted conservation can be problematic (Kardos & Luikart, 2020; Shafer et al., 2015; Waples & Lindley, 2018), there are scenarios where large effect loci can be expected to provide direct applications for conservation management.

## Acknowledgements

We thank Meri Lindqvist, Ville Aukee, Yann Czorlich, Victoria Pritchard, Jan Laine, Ryan Bare, Samuel May, Isadora Jimenez-Hidalgo, Eleni Petrou, Carolyn Tarpey, Cassondra Columbus, Ellen Campbell, Elena Correa, Vanessa Morman, and Steven Micheletti for samples, project development, and laboratory assistance. This work was supported by the Academy of Finland (project numbers 284941, 286334, 314254, and 314255), the European Research Council under the European Articles Union’s Horizon 2020 research and innovation program (grant agreement number 742312), the University of Helsinki, Washington Sea Grant (award numbers R/HCE-4 and R/E/I-26), and the School of Aquatic and Fishery Sciences at the University of Washington. The findings and conclusions in this paper are those of the authors and do not necessarily represent the views of the National Marine Fisheries Service.

## Data Archiving Statement

The data that support the findings of this study will be openly available upon acceptance to a peer-reviewed journal.

## Supporting Information

**File S1**: Additional Supporting Information regarding population descriptions, principal component analyses to account for population structure, and obtaining molecular estimates of pairwise relatedness within populations.

**File S2**: Supplementary Tables S1-S17, including genotypes for target SNPs within six6 and vgll3 and all putatively neutral loci, genome alignments, primer information, and model results.

## Literature Cited

Ayala, F. J., & Campbell, C. A. (1974). Frequency-dependent selection. Annual Review of Ecology and Systematics, 5, 115–138.

Ayllon, F., Kjaerner-Semb, E., Furmanek, T., Wennevik, V., Solberg, M. F., Dahle, G., … Wargelius, A. (2015). The vgll3 locus controls age at maturity in wild and domesticated Atlantic salmon (Salmo salar L.) males. Plos Genetics, 11(11), 15. doi: 10.1371/journal.pgen.1005628

Ayllon, F., Solberg, M. F., Glover, K. A., Mohammadi, F., Kjaerner-Semb, E., Fjelldal, P. G., … Wargelius, A. (2019). The influence of vgll3 genotypes on sea age at maturity is altered in farmed mowi strain Atlantic salmon. Bmc Genetics, 20,8. doi: 10.1186/s12863-019-0745-9

Baetscher, D. S., Clemento, A. J., Ng, T. C., Anderson, E. C., & Garza, J. C. (2018). Microhaplotypes provide increased power from short-read DNA sequences for relationship inference. Molecular Ecology Resources, 18(2), 296–305. doi: 10.1111/1755-0998.12737

Barson, N. J., Aykanat, T., Hindar, K., Baranski, M., Bolstad, G. H., Fiske, P., … Primmer, C. R. (2015). Sex-dependent dominance at a single locus maintains variation in age at maturity in salmon. Nature, 528(7582), 405–408. doi: 10.1038/nature16062

Barton, K. (2016). Package ‘MuMIn.’ CRAN Repos at https://cran.r-project.org/web/packages/MuMIn/MuMIn.pdf.

Barton, N. H., & Keightley, P. D. (2002). Understanding quantitative genetic variation. Nature Reviews Genetics, 3(1), 11–21. doi: 10.1038/nrg700

Behnke, R. J. (2002). Trout and Salmon of North America. New York: Free Press.

Boulding, E. G., Ang, K. P., Elliott, J. A. K., Powell, F., & Schaeffer, L. R. (2019). Differences in genetic architecture between continents at a major locus previously associated with sea age at sexual maturity in European Atlantic salmon. Aquaculture, 500, 670–678. doi: 10.1016/j.aquaculture.2018.09.025

Bowersox, B. J., Corsi, M. P., McCormick, J. L., Copeland, T., & Campbell, M. R. (2019). Examining life-history shifts and genetic composition in a hatchery steelhead population, with implications for fishery and ocean selection. Transactions of the American Fisheries Society, 148(6), 1056–1068. doi: 10.1002/tafs.10199

Brown, C. L., & Godduhn, A. (2015). Socioeconomic effects of declining salmon runs on the Yukon River. (398). Fairbanks.

Campbell, N. R., Harmon, S. A., & Narum, S. R. (2015). Genotyping-in-Thousands by sequencing (GT-seq): A cost effective SNP genotyping method based on custom amplicon sequencing. Molecular Ecology Resources, 15(4), 855–867. doi: 10.1111/1755-0998.12357

Cánovas, A., Reverter, A., DeAtley, K. L., Ashley, R. L., Colgrave, M. L., Fortes, M. R. S., … Thomas, M. G. (2014). Multi-tissue omics analyses reveal molecular regulatory networks for puberty in composite beef cattle. Plos One, 9(7), 17. doi: 10.1371/journal.pone.0102551

Carlson, S. M., & Seamons, T. R. (2008). A review of quantitative genetic components of fitness in salmonids: implications for adaptation to future change. Evolutionary Applications, 1(2), 222–238. doi: 10.1111/j.1752-4571.2008.00025.x

Cauwelier, E., Gilbey, J., Sampayo, J., Stradmeyer, L., & Middlemas, S. J. (2018). Identification of a single genomic region associated with seasonal river return timing in adult Scottish Atlantic salmon (Salmo salar), using a genome-wide association study. Canadian Journal of Fisheries and Aquatic Sciences, 75(9), 1427–1435. doi: 10.1139/cjfas-2017-0293

Charlesworth, B., Nordborg, M., & Charlesworth, D. (1997). The effects of local selection, balanced polymorphism and background selection on equilibrium patterns of genetic diversity in subdivided populations. Genetics Research, 70(2), 155–174. doi: 10.1017/s0016672397002954

Charnov, E. L., & Gillooly, J. F. (2004). Size and temperature in the evolution of fish life histories. Integrative and Comparative Biology, 44(6), 494–497. doi: 10.1093/icb/44.6.494

Christensen, K. A., Gutierrez, A. P., Lubieniecki, K. P., & Davidson, W. S. (2017). TEAD3, implicated by association to grilsing in Atlantic salmon. Aquaculture, 479, 571–578. doi: 10.1016/j.aquaculture.2017.06.026

Christensen, R. H. B. (2019). ordinal-Regression models for ordinal data. R package version 2019. 12–10. https://CRAN.R-project.org/package=ordinal.

Cingolani, P., Platts, A., Wang, L. L., Coon, M., Nguyen, T., Wang, L., … Ruden, D. M. (2012). A program for annotating and predicting the effects of single nucleotide polymorphisms, SnpEff: SNPs in the genome of Drosophila melanogaster strain w(1118); iso-2; iso-3. Fly, 6(2), 80–92. doi: 10.4161/fly.19695

Clark, D. D., Gorman, M. R., Hatori, M., Meadows, J. D., Panda, S., & Mellon, P. L. (2013). Aberrant development of the suprachiasmatic nucleus and circadian rhythms in mice lacking the homeodomain protein six6. Journal of Biological Rhythms, 28(1), 15–25. doi: 10.1177/0748730412468084

Cline, T. J., Ohlberger, J., & Schindler, D. E. (2019). Effects of warming climate and competition in the ocean for life-histories of Pacific salmon. Nature Ecology & Evolution, 3(6), 935–942. doi: 10.1038/s41559-019-0901-7

Conover, D. O., & Munch, S. B. (2002). Sustaining fisheries yields over evolutionary time scales. Science, 297(5578), 94–96. doi: 10.1126/science.1074085

Copeland, T., Ackerman, M. W., Wright, K. K., & Byrne, A. (2017). Life-history diversity of Snake River steelhead populations between and within management categories. North American Journal of Fisheries Management, 37(2), 395–404. doi: 10.1080/02755947.2016.1264506

Cortez, D., Marin, R., Toledo-Flores, D., Froidevaux, L., Liechti, A., Waters, P. D., … Kaessmann, H. (2014). Origins and functional evolution of Y chromosomes across mammals. Nature, 508(7497), 488-+. doi: 10.1038/nature13151

Cousminer, D. L., Berry, D. J., Timpson, N. J., Ang, W., Thiering, E., Byrne, E. M., … Early Growth Genetics, E. G. G. (2013). Genome-wide association and longitudinal analyses reveal genetic loci linking pubertal height growth, pubertal timing and childhood adiposity. Human Molecular Genetics, 22(13), 2735–2747. doi: 10.1093/hmg/ddt104

Cousminer, D. L., Widen, E., & Palmert, M. R. (2016). The genetics of pubertal timing in the general population: recent advances and evidence for sex-specificity. Current Opinion in Endocrinology Diabetes and Obesity, 23(1), 57–65. doi: 10.1097/med.0000000000000213

Crete-Lafreniere, A., Weir, L. K., & Bernatchez, L. (2012) Framing the Salmonidae Family Phylogenetic Portrait: A More Complete Picture from Increased Taxon Sampling. Plos One, 7(10), 19. doi: 10.1371/journal.pone.0046662

Czorlich, Y., Aykanat, T., Erkinaro, J., Orell, P., & Primmer, C. R. (2018). Rapid sex-specific evolution of age at maturity is shaped by genetic architecture in Atlantic salmon. Nature Ecology & Evolution, 2(11), 1800–1807. doi: 10.1038/s41559-018-0681-5

Debes, P. V., Piavchenko, N., Ruokolainen, A., Ovaskainen, O., Moustakas-Verho, J. E., Parre, N., … Primmer, C. R. (2019). Large single-locus effects for maturation timing are mediated via condition variation in Atlantic salmon. bioRxiv. doi: https://doi.org/10.1101/780437

DeFilippo, L. B., Schindler, D. E., Ohlberger, J., Schaberg, K. L., Foster, M. B., Ruhl, D., & Punt, A. E. (2019). Recruitment variation disrupts the stability of alternative life histories in an exploited salmon population. Evolutionary Applications, 12(2), 214–229. doi: 10.1111/eva.12709

Devlin, B., & Roeder, K. (1999). Genomic control for association studies. Biometrics, 55(4), 997–1004. doi: 10.1111/j.0006-341X.1999.00997.x

Elmer, K. R., & Meyer, A. (2011). Adaptation in the age of ecological genomics: insights from parallelism and convergence. Trends in Ecology & Evolution, 26(6), 298–306. doi: 10.1016/j.tree.2011.02.008

Fariello, M. I., Boitard, S., Mercier, S., Robelin, D., Faraut, T., Arnould, C., … SanCristobal, M. (2017). Accounting for linkage disequilibrium in genome scans for selection without individual genotypes: The local score approach. Molecular Ecology, 26(14), 3700–3714. doi: 10.1111/mec.14141

Faust, G. G., & Hall, I. M. (2014). SAMBLASTER: fast duplicate marking and structural variant read extraction. Bioinformatics, 30(17), 2503–2505. doi: 10.1093/bioinformatics/btu314

Flatt, T., & Heyland, A. (2011). Mechanisms of Life-history Evolution: the genetics and physiology of life-history traits and trade-offs: Oxford University Press.

Fleming, I. A. (1996). Reproductive strategies of Atlantic salmon: Ecology and evolution. Reviews in Fish Biology and Fisheries, 6(4), 379–416. doi: 10.1007/bf00164323

Fraser, D. J., Weir, L. K., Bernatchez, L., Hansen, M. M., & Taylor, E. B. (2011). Extent and scale of local adaptation in salmonid fishes: review and meta-analysis. Heredity, 106(3), 404–420. doi: 10.1038/hdy.2010.167

Freese, N. H., Norris, D. C., & Loraine, A. E. (2016). Integrated genome browser: visual analytics platform for genomics. Bioinformatics, 32(14), 2089–2095. doi: 10.1093/bioinformatics/btw069

Giska, I., Farelo, L., Pimenta, J., Seixas, F. A., Ferreira, M. S., Marques, J. P., … Melo-Ferreira, J. (2019). Introgression drives repeated evolution of winter coat color polymorphism in hares. Proceedings of the National Academy of Sciences of the United States of America, 116(48), 24150–24156. doi: 10.1073/pnas.1910471116

Gjerde, B. (1984). Response to individual selection for age at sexual maturity in Atlantic salmon. Aquaculture, 38(3), 229–240. doi: 10.1016/0044-8486(84)90147-9

Greene, C. M., Hall, J. E., Guilbault, K. R., & Quinn, T. P. (2010). Improved viability of populations with diverse life-history portfolios. Biology Letters, 6(3), 382–386. doi: 10.1098/rsbl.2009.0780

Guerrero, R. F., & Hahn, M. W. (2017). Speciation as a sieve for ancestral polymorphism. Molecular Ecology, 26(20), 5362–5368. doi: 10.1111/mec.14290

Hankin, D. G., & Healey, M. C. (1986) Dependence of exploitation rates for maximum yield and stock collapse on age and sex structure of Chinook salmon (Oncorhynchus tshawytscha) stocks. Canadian Journal of Fisheries and Aquatic Sciences, 43(9), 1746–1759. doi: 10.1139/f86-219

Hankin, D. G., Nicholas, J. W., & Downey, T. W. (1993). Evidence for inheritance of age at maturity in Chinook salmon (Oncorhynchus tshawytscha). Canadian Journal of Fisheries and Aquatic Sciences, 50(2), 347–358. doi: 10.1139/f93-040

Harstad, D. L., Larsen, D. A., & Beckman, B. R. (2014). Variation in minijack rate among hatchery populations of Columbia river basin Chinook salmon. Transactions of the American Fisheries Society, 143(3), 768–778. doi: 10.1080/00028487.2014.886621

Harstad, D. L., Larsen, D. A., Miller, J., Adams, I., Spangenberg, D. K., Nance, S., … Beckman, B. R. (2018). Winter-rearing temperature affects growth profiles, age of maturation, and smolt-to-adult returns for yearling summer Chinook salmon in the Upper Columbia River Basin. North American Journal of Fisheries Management, 38(4), 867–885. doi: 10.1002/nafm.10186

Healey, M. C. (1991). Life-history in Chinook salmon (Oncorhynchus tshawytscha). In C. Groot & L. Margolis (Eds.), Pacific salmon life histories (pp. 311-393). Vancouver, Canada: University of British Columbia Press.

Hess, J. E., Ackerman, M. W., Fryer, J. K., Hasselman, D. J., Steele, C. A., Stephenson, J. J., … Narum, S. R. (2016). Differential adult migration-timing and stock-specific abundance of steelhead in mixed stock assemblages. Ices Journal of Marine Science, 73(10), 2606–2615. doi: 10.1093/icesjms/fsw138

Hou, H. Y., Uuskula-Reimand, L., Makarem, M., Corre, C., Saleh, S., Metcalf, A., … Wilson, M. D. (2017). Gene expression profiling of puberty-associated genes reveals abundant tissue and sex-specific changes across postnatal development. Human Molecular Genetics, 26(18), 3585–3599. doi: 10.1093/hmg/ddx246

Jamie, G. A., & Meier, J. I. (2020). The persistence of polymorphisms across species radiations. Trends in Ecology and Evolution. doi: 10.1016/j.tree.2020.04.007

Janowitz-Koch, I., Rabe, C., Kinzer, R., Nelson, D., Hess, M. A., & Narum, S. R. (2019). Long-term evaluation of fitness and demographic effects of a Chinook Salmon supplementation program. Evolutionary Applications, 12(3), 456–469. doi: 10.1111/eva.12725

Jean, D., Bernier, G., & Gruss, P. (1999). Six6 (Optx2) is a novel murine Six3-related homeobox gene that demarcates the presumptive pituitary/hypothalamic axis and the ventral optic stalk. Mechanisms of Development, 84(1-2), 31–40. doi: 10.1016/s0925-4773(99)00068-4

Johnston, S. E., Orell, P., Pritchard, V. L., Kent, M. P., Lien, S., Niemela, E., … Primmer, C. R. (2014). Genome-wide SNP analysis reveals a genetic basis for sea-age variation in a wild population of Atlantic salmon (Salmo salar). Molecular Ecology, 23(14), 3452–3468. doi: 10.1111/mec.12832

Jombart, T. (2008). adegenet: a R package for the multivariate analysis of genetic markers. Bioinformatics, 24(11), 1403–1405. doi: 10.1093/bioinformatics/btn129

Jombart, T., & Ahmed, I. (2011). adegenet 1.3-1: new tools for the analysis of genome-wide SNP data. Bioinformatics, 27(21), 3070–3071. doi: 10.1093/bioinformatics/btr521

Jones, F. C., Grabherr, M. G., Chan, Y. F., Russell, P., Mauceli, E., Johnson, J., … Whole Genome Assembly, T. (2012). The genomic basis of adaptive evolution in threespine sticklebacks. Nature, 484(7392), 55–61. doi: 10.1038/nature10944

Kardos, M., & Luikart, G. (2020). The genetic architecture of fitness drives population viability during rapid environmental change. BioRxiv:660803.

Keefer, M. L., & Caudill, C. C. (2014). Homing and straying by anadromous salmonids: a review of mechanisms and rates. Reviews in Fish Biology and Fisheries, 24(1), 333–368. doi: 10.1007/s11160-013-9334-6

Kenny, D. A., Heslin, J., & Byrne, C. J. (2018). Early onset of puberty in cattle: implications for gamete quality and embryo survival. Reproduction Fertility and Development, 30(1), 101–117. doi: 10.1071/rd17376

Kindsvater, H. K., Mangel, M., Reynolds, J. D., & Dulvy, N. K. (2016). Ten principles from evolutionary ecology essential for effective marine conservation. Ecology and Evolution, 6(7), 2125–2138. doi: 10.1002/ece3.2012

Kjaerner-Semb, E., Ayllon, F., Kleppe, L., Sorhus, E., Skaftnesmo, K., Furmanek, T., … Edvardsen, R. B. (2018). Vgll3 and the Hippo pathway are regulated in Sertoli cells upon entry and during puberty in Atlantic salmon testis. Scientific Reports, 8, 11. doi: 10.1038/s41598-018-20308-1

Kodama, M., Brieuc, M. S. O., Devlin, R. H., Hard, J. J., & Naish, K. A. (2014). Comparative mapping Between coho salmon (Oncorhynchus kisutch) and three other salmonids suggests a role for chromosomal rearrangements in the retention of duplicated regions following a whole genome duplication event. G3-Genes Genomes Genetics, 4(9), 1717–1730. doi: 10.1534/g3.114.012294

Kodama, M., Hard, J. J., & Naish, K. A. (2018). Mapping of quantitative trait loci for temporal growth and age at maturity in coho salmon: Evidence for genotype-by-sex interactions. Marine Genomics, 38, 33–44. doi: 10.1016/j.margen.2017.07.004

Kofler, R., Pandey, R. V., & Schlotterer, C. (2011). PoPoolation2: identifying differentiation between populations using sequencing of pooled DNA samples (Pool-Seq). Bioinformatics, 27(24), 3435–3436. doi: 10.1093/bioinformatics/btr589

Koressaar, T., & Remm, M. (2007). Enhancements and modifications of primer design program Primer3. Bioinformatics, 23(10), 1289–1291. doi: 10.1093/bioinformatics/btm091

Korte, A., & Farlow, A. (2013). The advantages and limitations of trait analysis with GWAS: a review. Plant Methods, 9, 9. doi: 10.1186/1746-4811-9-29

Kozlowski, J. (1992). Optimal allocation of resources to growth and reproduction - implications for age and size at maturity. Trends in Ecology & Evolution, 7(1), 15–19. doi: 10.1016/0169-5347(92)90192-e

Kronforst, M. R., Barsh, G. S., Kopp, A., Mallet, J., Monteiro, A., Mullen, S. P., … Hoekstra, H. E. (2012). Unraveling the thread of nature’s tapestry: the genetics of diversity and convergence in animal pigmentation. Pigment Cell & Melanoma Research, 25(4), 411–433. doi: 10.1111/j.1755-148X.2012.01014.x

Kumar, J. P. (2009). The sine oculis homeobox (SIX) family of transcription factors as regulators of development and disease. Cellular and Molecular Life Sciences, 66(4), 565–583. doi: 10.1007/s00018-008-8335-4

Kurko, J., Debes, P. V., House, A. H., Aykanat, T., Erkinaro, J., & Primmer, C. R. (2020). Transcription profiles of age-at-maturity-associated genes suggest cell fate commitment regulation as a key factor in the Atlantic salmon maturation process. G3-Genes Genomes Genetics, 10(1), 235–246. doi: 10.1534/g3.119.400882

Langmead, B., & Salzberg, S. L. (2012). Fast gapped-read alignment with Bowtie 2. Nature Methods, 9(4), 357–U354. doi: 10.1038/nmeth.1923

Larder, R., Clark, D. D., Miller, N. L. G., & Mellon, P. L. (2011). Hypothalamic dysregulation and infertility in mice lacking the homeodomain protein six6. Journal of Neuroscience, 31(2), 426–438. doi: 10.1523/jneurosci.1688-10.2011

Larsen, D. A., Harstad, D. L., Fuhrman, A. E., Knudsen, C. M., Schroder, S. L., Bosch, W. J., … Beckman, B. R. (2019). Maintaining a wild phenotype in a conservation hatchery program for Chinook salmon: The effect of managed breeding on early male maturation. Plos One, 14(5), 32. doi: 10.1371/journal.pone.0216168

Lewis, B., Grant, W. S., Brenner, R. E., & Hamazaki, T. (2015). Changes in size and age of Chinook salmon Oncorhynchus tshawytscha returning to Alaska. Plos One, 10(6), 17. doi: 10.1371/journal.pone.0130184

Li, C. Y., Kan, L. J., Chen, Y., Zheng, X. D., Li, W. N., Zhang, W. X., … Chen, D. H. (2015). Ci antagonizes Hippo signaling in the somatic cells of the ovary to drive germline stem cell differentiation. Cell Research, 25(10), 1152–1170. doi: 10.1038/cr.2015.114

Li, H. (2011). A statistical framework for SNP calling, mutation discovery, association mapping and population genetical parameter estimation from sequencing data. Bioinformatics, 27(21), 2987–2993. doi: 10.1093/bioinformatics/btr509

Li, H. (2013). Aligning sequence reads, clone sequences and assembly contigs with BWA-MEM. arXiv. doi: 1303.3997v2

Li, H., Handsaker, B., Wysoker, A., Fennell, T., Ruan, J., Homer, N., … Genome Project Data, P. (2009). The sequence alignment/map format and SAMtools. Bioinformatics, 25(16), 2078–2079. doi: 10.1093/bioinformatics/btp352

Lien, S., Koop, B. F., Sandve, S. R., Miller, J. R., Kent, M. P., Nome, T., … Davidson, W. S. (2016). The Atlantic salmon genome provides insights into rediploidization. Nature, 533(7602), 200-+. doi: 10.1038/nature17164

Liu, S. L., Ferchaud, A. L., Gronkjaer, P., Nygaard, R., & Hansen, M. M. (2018). Genomic parallelism and lack thereof in contrasting systems of three-spined sticklebacks. Molecular Ecology, 27(23), 4725–4743. doi: 10.1111/mec.14782

Macqueen, D. J., & Johnston, I. A. (2014). A well-constrained estimate for the timing of the salmonid whole genome duplication reveals major decoupling from species diversification. Proceedings of the Royal Society B-Biological Sciences, 281(1778), 8. doi: 10.1098/rspb.2013.2881

Mank, J. E. (2017). Population genetics of sexual conflict in the genomic era. Nature Reviews Genetics, 18(12), 721–730. doi: 10.1038/nrg.2017.83

Mank, J. E., & Avise, J. C. (2006). The evolution of reproductive and genomic diversity in ray-finned fishes: insights from phylogeny and comparative analysis. Journal of Fish Biology, 69(1), 1–27. doi: 10.1111/j.1095-8649.2006.01132.x

Marchini, J., Cardon, L. R., Phillips, M. S., & Donnelly, P. (2004). The effects of human population structure on large genetic association studies. Nature Genetics, 36(5), 512–517. doi: 10.1038/ng1337

McKinney, G. J., Nichols, K. M., & Ford, M. J. (2020). A mobile sex-determining region, male-specific haplotypes, and rearing environment influence age at maturity in Chinook salmon. bioRxiv. doi: https://doi.org/10.1101/2020.04.23.056093

McKinney, G. J., Seeb, J. E., Pascal, C. E., Schindler, D. E., Gilk-Baumer, S. E., & Seeb, L. W. (2019). Y-chromosome haplotypes drive variation in size and age at maturity in male Chinook salmon. bioRxiv. doi: https://doi.org/10.1101/691063

Micheletti, S. J., Hess, J. E., Zendt, J. S., & Narum, S. R. (2018). Selection at a genomic region of major effect is responsible for evolution of complex life histories in anadromous steelhead. Bmc Evolutionary Biology, 18, 11. doi: 10.1186/s12862-018-1255-5

Micheletti, S. J., & Narum, S. R. (2018). Utility of pooled sequencing for association mapping in nonmodel organisms. Molecular Ecology Resources, 18(4), 825–837. doi: 10.1111/1755-0998.12784

Mobley, K. B., Granroth-Wilding, H., Ellmen, M., Vaha, J. P., Aykanat, T., Johnston, S. E., … Primmer, C. R. (2019). Home ground advantage: Local Atlantic salmon have higher reproductive fitness than dispersers in the wild. Science Advances, 5(2), 8. doi: 10.1126/sciadv.aav1112

Mohamed, A. R., Verbyla, K. L., Al-Mamun, H. A., McWilliam, S., Evans, B., King, H., … Kijas, J. W. (2019). Polygenic and sex specific architecture for two maturation traits in farmed Atlantic salmon. Bmc Genomics, 20, 12. doi: 10.1186/s12864-019-5525-4

Mundy, N. I. (2005). A window on the genetics of evolution: MC1R and plumage colouration in birds. Proceedings of the Royal Society B-Biological Sciences, 272(1573), 1633–1640. doi: 10.1098/rspb.2005.3107

Nielsen, J. L. (1999). The evolutionary history of steelhead (Oncorhynchus mykiss) along the US Pacific Coast: Developing a conservation strategy using genetic diversity. Ices Journal of Marine Science, 56(4), 449–458. doi: 10.1006/jmsc.1999.0452

Ohlberger, J., Schindler, D. E., Brown, R. J., Harding, J. M. S., Adkison, M. D., Munro, A. R., … Spaeder, J. (2020). The reproductive value of large females: consequences of shifts in demographic structure for population reproductive potential in Chinook salmon. Canadian Journal of Fisheries and Aquatic Sciences.

Ohlberger, J., Ward, E. J., Schindler, D. E., & Lewis, B. (2018). Demographic changes in Chinook salmon across the Northeast Pacific Ocean. Fish and Fisheries, 19(3), 533–546. doi: 10.1111/faf.12272

Oomen, R. A., Kuparinen, A., & Hutchings, J. A. (2020). Consequences of single-locus and tightly linked genomic architectures for evolutionary responses to environmental change. Journal of Heredity.

Pearse, D. E., Barson, N. J., Nome, T., Gao, G. T., Campbell, M. A., Abadia-Cardoso, A., … Lien, S. (2019). Sex-dependent dominance maintains migration supergene in rainbow trout. Nature Ecology & Evolution, 3(12), 1731-+. doi: 10.1038/s41559-019-1044-6

Perry, J. R. B., Day, F., Elks, C. E., Sulem, P., Thompson, D. J., Ferreira, T., … Early Growth Genetics, E. G. G. C. (2014). Parent-of-origin-specific allelic associations among 106 genomic loci for age at menarche. Nature, 514(7520), 92-+. doi: 10.1038/nature13545

Price, A. L., Patterson, N. J., Plenge, R. M., Weinblatt, M. E., Shadick, N. A., & Reich, D. (2006). Principal components analysis corrects for stratification in genome-wide association studies. Nature Genetics, 38(8), 904–909. doi: 10.1038/ng1847

Pritchard, V. L., Makinen, H., Vaha, J. P., Erkinaro, J., Orell, P., & Primmer, C. R. (2018). Genomic signatures of fine-scale local selection in Atlantic salmon suggest involvement of sexual maturation, energy homeostasis and immune defence-related genes. Molecular Ecology, 27(11), 2560–2575. doi: 10.1111/mec.14705

Quinn, T. P. (2005). The Behavior and Ecology of Pacific Salmon and Trout. Seattle: University of Washington Press.

Quinn, T. P., Wetzel, L., Bishop, S., Overberg, K., & Rogers, D. E. (2001). Influence of breeding habitat on bear predation and age at maturity and sexual dimorphism of sockeye salmon populations. Canadian Journal of Zoology, 79(10), 1782–1793. doi: 10.1139/cjz-79-10-1782

R_Core_Team. (2019). R: A language and environment for statistical computing. Vienna, Austria: R Foundation for Statistical Computing. http://www.R-project.org. Retrieved from http://www.R-project.org

Ricker, W. E. (1980). Causes of the decrease in age and size of Chinook salmon (Oncorhynchus tshawytscha). (944). Nanaimo, British Columbia.

Roff, D. A. (2011). Alternative strategies: The evolution of switch points. Current Biology, 21(8), R285–R287. doi: 10.1016/j.cub.2011.03.016

Roff, D. A., Mostowy, S., & Fairbairn, D. J. (2002). The evolution of trade-offs: Testing predictions on response to selection and environmental variation. Evolution, 56(1), 84–95.

Roni, P., & Quinn, T. P. (1995). Geographic variation in size and age of North American Chinook salmon. North American Journal of Fisheries Management, 15(2), 325–345.

Samy, J. K. A., Mulugeta, T. D., Nome, T., Sandve, S. R., Grammes, F., Kent, M. P., … Vage, D. I. (2017). SalmoBase: an integrated molecular data resource for Salmonid species. Bmc Genomics, 18, 5. doi: 10.1186/s12864-017-3877-1

Satterthwaite, W. H., Carlson, S. M., & Criss, A. (2017). Ocean size and corresponding life-history diversity among the four run timings of California Central Valley Chinook salmon. Transactions of the American Fisheries Society, 146(4), 594–610. doi: 10.1080/00028487.2017.1293562

Schaffer, W. M. (2004). Life histories, evolution, and salmonids. In A. P. Hendry & S. C. Stearns (Eds.), Evolution Illuminated: Salmon and Their Relatives: Oxford University Press.

Schindler, D. E., Hilborn, R., Chasco, B., Boatright, C. P., Quinn, T. P., Rogers, L. A., & Webster, M. S. (2010). Population diversity and the portfolio effect in an exploited species. Nature, 465(7298), 609–612. doi: 10.1038/nature09060

Schlötterer, C., Tobler, R., Kofler, R., & Nolte, V. (2014). Sequencing pools of individuals-mining genome-wide polymorphism data without big funding. Nature Reviews Genetics, 15(11), 749–763. doi: 10.1038/nrg3803

Schwartz, M. K., Luikart, G., & Waples, R. S. (2007). Genetic monitoring as a promising tool for conservation and management. Trends in Ecology & Evolution, 22(1), 25–33. doi: 10.1016/j.tree.2006.08.009

Seo, H. C., Drivenes, O., Ellingsen, S., & Fjose, A. (1998). Expression of two zebrafish homologues of the murine Six3 gene demarcates the initial eye primordia. Mechanisms of Development, 73(1), 45–57. doi: 10.1016/s0925-4773(98)00028-8

Shafer, A. B. A., Wolf, J. B. W., Alves, P. C., Bergstrom, L., Bruford, M. W., Brannstrom, I., … Zielinski, P. (2015). Genomics and the challenging translation into conservation practice. Trends in Ecology & Evolution, 30(2), 78–87. doi: 10.1016/j.tree.2014.11.009

Simon, E., Faucheux, C., Zider, A., Theze, N., & Thiebaud, P. (2016). From vestigial to vestigial-like: the Drosophila gene that has taken wing. Development Genes and Evolution, 226(4), 297–315. doi: 10.1007/s00427-016-0546-3

Sinclair-Waters, M., Odegard, J., Korsvoll, S. A., Moen, T., Lien, S., Primmer, C. R., & Barson, N. J. (2020). Beyond large-effect loci: large-scale GWAS reveals a mixed large-effect and polygenic architecture for age at maturity of Atlantic salmon. Genetics Selection Evolution, 52(1), 11. doi: 10.1186/s12711-020-0529-8

Stearns, S. C. (1989). Trade-offs in life-history evolution. Functional Ecology, 3(3), 259–268. doi: 10.2307/2389364

Stearns, S. C. (1992). The Evolution of Life Histories. London: Oxford University Press.

Taranger, G. L., Carrillo, M., Schulz, R. W., Fontaine, P., Zanuy, S., Felip, A., … Hausen, T. (2010). Control of puberty in farmed fish. General and Comparative Endocrinology, 165(3), 483–515. doi: 10.1016/j.ygcen.2009.05.004

Taylor, E. B. (1990). Environmental correlates of life-history variation in juvenile Chinook salmon, Oncorhynchus tshawytscha (Walbaum). Journal of Fish Biology, 37(1), 1–17. doi: 10.1111/j.1095-8649.1990.tb05922.x

Taylor, E. B. (1991). A review of local adaptation in Salmonidae, with particular reference to Pacific and Atlantic salmon. Aquaculture, 98(1-3), 185–207. doi: 10.1016/0044-8486(91)90383-i

Tipping, J. M. (1991). Heritability of age at maturity in steelhead. North American Journal of Fisheries Management, 11, 105–108. doi: https://doi.org/10.1577/1548-8675(1991)011<0105:MBHOAA>2.3.CO;2

Turner, S. D. (2014). qqman: an R package for visualizing GWAS results using Q-Q and manhattan plots. bioRxiv. doi: https://doi.org/10.1101/005165

Untergasser, A., Cutcutache, I., Koressaar, T., Ye, J., Faircloth, B. C., Remm, M., & Rozen, S. G. (2012). Primer3-new capabilities and interfaces. Nucleic Acids Research, 40(15), 12. doi: 10.1093/nar/gks596

Vähä, J. P., Erkinaro, J., Niemela, E., & Primmer, C. R. (2008). Temporally stable genetic structure and low migration in an Atlantic salmon population complex: implications for conservation and management. Evolutionary Applications, 1(1), 137–154. doi: 10.1111/j.1752-4571.2007.00007.x

Veale, A. J., & Russello, M. A. (2017). An ancient selective sweep linked to reproductive life-history evolution in sockeye salmon. Scientific Reports, 7, 10. doi: 10.1038/s41598-017-01890-2

Verta, J.-P., Debes, P. V., Piavchenko, N., Ruokolainen, A., Ovaskainen, O., Moustakas-Verho, J. E., … Primmer, C. R. (2019). Regulatory divergence in vgll3 underlies variation in age at maturity in male Atlantic salmon. bioRxiv. doi: https://doi.org/10.1101/777300

Waples, R. S., & Lindley, S. T. (2018). Genomics and conservation units: The genetic basis of adult migration timing in Pacific salmonids. Evolutionary Applications, 11(9), 1518–1526. doi: 10.1111/eva.12687

Waples, R. S., Teel, D. J., Myers, J. M., & Marshall, A. R. (2004). Life-history divergence in Chinook salmon: Historic contingency and parallel evolution. Evolution, 58(2), 386–403. doi: 10.1111/j.0014-3820.2004.tb01654.x

Watanabe, N., Itoh, K., Mogi, M., Fujinami, Y., Shimizu, D., Hashimoto, H., … Suzuki, T. (2012). Circadian pacemaker in the suprachiasmatic nuclei of teleost fish revealed by rhythmic period2 expression. General and Comparative Endocrinology, 178(2), 400–407. doi: 10.1016/j.ygcen.2012.06.012

Yano, A., Nicol, B., Jouanno, E., Quillet, E., Fostier, A., Guyomard, R., & Guiguen, Y. (2013). The sexually dimorphic on the Y-chromosome gene (sdY) is a conserved male-specific Y-chromosome sequence in many salmonids. Evolutionary Applications, 6(3), 486–496. doi: 10.1111/eva.12032

Yeaman, S., & Whitlock, M. C. (2011). The genetic architecture of adaptation under migration-selection balance. Evolution, 65(7), 1897–1911. doi: 10.1111/j.1558-5646.2011.01269.x

