## Supplementary material for "Heterogeneous genetic basis of age at maturity in salmonid fishes": File S1 Supplementary Information

*Sample collections*

*Chinook salmon*

**Cle Elum:** The Cle Elum Supplementation and Research Facility (CESRF) in Cle Elum, Washington, U.S.A. (47°18'N, 120°96'W) was founded from 1997–2002 using returning wild adults from the upper Yakima River population. In 2002, the hatchery population was divided into an integrated (INT) line and a segregated (SEG) line by spawning wild ($F_1$ Wild) and first generation hatchery-origin ($F_1$ Hatchery) adults, respectively (Waters et al., 2018; Waters et al., 2015). Broodstock for the integrated line comprise only fish born in the wild, and all returning adults from this line are allowed to spawn in the river. The segregated line, in contrast, uses only returning hatchery-origin fish as broodstock, and no adults are allowed to reproduce naturally. Juveniles typically spend approximately 18 months in freshwater before being released to begin their migration to the ocean. A majority (>75%, Knudsen et al., 2006) of Chinook from CESRF spend two years in the ocean and return at age four to reproduce. However, males may mature at two to five years of age while females may mature at three to five years. Ages at maturity of hatchery-origin broodstock are determined from passive integrated transponder or coded wire tags, which denote the brood year of each fish, while those of natural-origin broodstock and hatchery-origin fish without tags were determined from growth rings on their scales (Clutter & Whitesel, 1956). Samples for this study were collected between 1998 and 2014.

**Feather River:** The Feather River is the primary tributary to the Sacramento River in California's northern Central Valley, maintaining both spring-run (enter freshwater in Spring, months prior to spawning) and fall-run (enter freshwater in Fall shortly before spawning) populations. Oroville Dam was constructed in 1967 as a principal feature of the California State Water Project, effectively eliminating the primary spawning habitat in the drainage, with disproportionate impacts on the spring-run population. The Feather River Hatchery (39°31'N, 121°33'W) was constructed as mitigation for that lost spawning habitat. Management of the hatchery led to widespread introgression between the spring-run and fall-run populations until 2006, when hatchery operations were modified to require observation of the spring-run phenotype (early return time) for inclusion in that population. Ongoing genetic monitoring of the spring-run hatchery population via parentage (Clemento, 2013) was used to calculate the ages used in this study. Samples were collected by hatchery staff during adult spawning from 2009 to 2017. Juveniles are reared at the hatchery until they achieve smolt size (<1 year old) and are then released to the river for direct outmigration.

**Little Port Walter:** NOAA's Little Port Walter Research Station (LPW), located on southern Baranof Island, Alaska, U.S.A. (56°38'N, 134°64'W), maintained two hatchery lines of Chinook salmon. The lines were established using wild gametes collected from the Chickamin and Unuk rivers, located approximately 250 km from LPW near Ketchikan. Wild gametes were collected from the Chickamin river in 1976 and again in 1996 (Templin, 2001). Wild gametes from the Unuk river were collected annually from 1976-1981 and again in 1998 (Templin, 2001). Juveniles are reared in freshwater for approximately 20 months before being released as outmigrating smolts. All smolts are coded wire tagged and have their adipose fins removed for external identification, which enable the stock and ages of all returning adults to be accurately determined. Adult males return at three to six years and females at four to six years. Samples for this study were collected from 2013 to 2016. The Chickamin hatchery line at LPW was discontinued in 2016.

**Sacramento River:** Sacramento winter-run Chinook salmon currently spawn in ~20km of the upper Sacramento River, below Keswick Dam (40°36'N, 122°26'W), the terminus of anadromy (Myers et al., 1998). The population was classified as endangered under the United States Endangered Species Act in 1994. Adults return to the river after one to four years in the ocean. Juveniles spend from 5-10 months feeding in freshwater before outmigrating as smolts. Samples were collected from returning adults at the Keswick Trap from 2012 to 2017, and ages were calculated using genetic-based parentage.

*Coho salmon*

**Big Beef Creek, Washington State:** Big Beef Creek is located in Hood Canal in Puget Sound, Washington State, U.S.A. (47°39'N, 122°46'W). There is no hatchery in this system; the population comprises naturally spawning individuals. Juveniles spend 18 months in freshwater, and adults returning to this system are either two years old (males only, jacks) or three years old (males and females). Returning adults were collected over two cohorts at a weir at the mouth of the creek during the years 2008 to 2009. Individuals were aged using molecular based pedigree analyses (Kodama, Hard, & Naish, 2012).

**Klamath River:** The Klamath River is the second largest river in California, U.S.A. and drains a large watershed that spans Northern California and south-central Oregon. Coho salmon in the Klamath River (part of the Southern Oregon/Northern California [SONCC] Evolutionary

Significant Unit) were listed as threatened under the United States Endangered Species Act in 1997. The Coho salmon population in the Klamath River is supplemented by production at Iron Gate Hatchery (41°55’N, 122°26’W) in the upper drainage. Juveniles are reared at the hatchery and released as outmigrating smolts at age-1+ after 16-18 months in freshwater. Samples for this study were collected from returning adults from 2009 to 2012, and ages were calculated using genetic pedigree reconstruction (Starks, 2014).

*Sockeye salmon*

**A and C Creeks, Wood River System, Alaska:** The A and C Creeks are located in the Little Togiak Lake, Wood River system, Bristol Bay, Alaska (A Creek: 59°34'N 159°07'W; C Creek: 59°34'N 159°09’W). The two small creeks (A - 350m long; C - 450m long) are less than 1km apart and support two wild populations that have been extensively sampled as part of a long-term field collection program (Lin, Quinn, Hilborn, & Hauser, 2008; May, McKinney, Hilborn, Hauser, & Naish, 2020; Peterson, Hilborn, & Hauser, 2014). Juveniles rear in streams and lakes for 18 months before migrating to the ocean. Adult males return at ages three to six while females return at ages four to six. However, the majority of the population comprises four and five year olds. Adults were sampled from 2008 to 2010. Age at maturity was determined directly using molecular pedigree analyses (Lin et al., 2008; Peterson et al., 2014).

*Steelhead trout*

**Dworshak:** Dworshak Hatchery (46°30'N, 116°19'W) is located in the Clearwater River in Idaho, U.S.A. and raises steelhead to mitigate for populations impacted by human activities including hydroelectric dams with limited fish passage. Steelhead produced at this hatchery are reared in freshwater for one year before release and are known to have a large proportion of older aged (2-ocean) fish than most other steelhead in the region, which typically spend only one year in the ocean (Bowersox, Corsi, McCormick, Copeland, & Campbell, 2019; Hess et al., 2016). Phenotypic variation for ocean-age is greater in males (predominantly 1-2 years in the ocean with rare cases of 3-4 years in the ocean) than females that are largely 2-ocean age. Samples for this study were collected in 2015 and 2016. The origin of each fish was verified through parentage analyses (see Hess et al., 2016 for details), ocean age at maturation was determined by examination of scale annuli collected from each fish, and total age was determined from genetic pedigrees. Body size was measured by fork length to the nearest centimeter and was expected to be directly correlated with years in the ocean for both sexes.

**Eel River:** The Eel River is the third largest river in California, draining the coastal mountains of Mendocino and Humboldt counties in Northern California. There is no hatchery in the system. The Eel River is designated Wild & Scenic at both the State (1972) and Federal (1981) levels. While the Wild & Scenic designation protects the river from construction of new dams and diversions, there is a legacy dam and fish counting station (Cape Horn Dam, constructed in 1907, 39°23’N, 123°07’W) that blocks access to the upper mainstem. As with many *O. mykiss* populations, some juvenile fish remain in freshwater as resident Rainbow Trout, while those that outmigrate as smolting Steelhead tend to do so after 16-30 months in freshwater. Samples for this study were collected from returning adult winter-run Steelhead between 2012 and 2015 before they were passed above the dam. Ages were determined via genetic pedigree reconstruction (unpublished data).

**Forks Creek, Washington State:** Forks Creek hatchery (46°33'N, 123°35’W) is located on a small tributary to the Willapa River in southwestern Washington State, U.S.A. In 1967, the hatchery initiated a winter line comprising native and introduced fish from Puget Sound, Washington (Crawford, 1979). Subsequent to 1994, the run was locally propagated. Juveniles are reared for 18 months in freshwater. Males mature at ages two to six years, while females mature at three to six years of age. However, the most frequent age class is three years. Returning adults were collected at a permanent weir spanning the Creek between 2000 and 2009. Ages at maturity were determined directly from molecular pedigree analysis (Naish, Seamons, Dauer, Hauser, & Quinn, 2013).

**Pahsimeroi:** Pahsimeroi Hatchery (44°41'N, 114°02'W) is located in the upper Salmon River in Idaho and raises steelhead for harvest mitigation purposes due to human impacts in the region that have negatively affected stocks. Steelhead produced at this hatchery are reared in freshwater for one year before release and typically spend only one year in the ocean with a low proportion of older aged (2-ocean) fish (Copeland, Ackerman, Wright, & Byrne, 2017; Hess et al., 2016). Phenotypic variation for ocean-age is greater in males (predominantly 1-ocean with cases of 2-ocean) than females that are largely 1-ocean age. Samples for this study were collected in 2015 and 2016. The origin of each fish was verified through parentage analyses (see Hess et al., 2016 for details), ocean age at maturation was determined by examination of scale annuli collected from each fish, and total age was determined from genetic pedigrees. Body size was measured by fork length to the nearest centimeter and was expected to be directly correlated with years in the ocean for both sexes.

**Wallowa:** Wallowa Hatchery (45°25'N, 117°18’W) is located in the Grande Ronde River in Oregon and raises steelhead for harvest mitigation purposes due to human impacts in the region that have negatively affected stocks. Steelhead produced at this hatchery are reared in freshwater for one year before release and typically spend only one year in the ocean with a low proportion of older aged (2-ocean) fish . Phenotypic variation for ocean-age is greater in males (predominantly 1-ocean with cases of 2-ocean) than females that are largely 1-ocean age. Samples for this study were collected in 2015 and 2016. The origin of each fish was verified through parentage analyses (see Hess et al., 2016 for details), ocean age at maturation was determined by examination of scale annuli collected from each fish, and total age was determined from genetic pedigrees. Body size was measured by fork length to the nearest centimeter and was expected to be directly correlated with years in the ocean for both sexes.

**Warm Springs:** Warm Springs Hatchery (38°43’N, 122°59’W) is located on Dry Creek, a tributary of the Russian River in central California, U.S.A. The hatchery opened in 1981 to mitigate the lost spawning and rearing habitat behind Warm Springs Dam. Hatchery produced fish are released at age 1+ as smolts (16-18 months) and return after 1-3 years at sea. Samples for this study were collected from returning adults from 2009 to 2011, and ages were calculated using genetic pedigree reconstruction (Abadía-Cardoso, Anderson, Pearse, & Garza, 2013).

*Principal component analyses to account for population structure*

A principal component analysis was conducted for each species using putatively neutral loci and the R package *adegenet* (v. 2.1.3, Jombart, 2008; Jombart & Ahmed, 2011). Principal components (PCs) that reflected population structure were identified using eigenvalues and the scree test (Cattell, 1966; D'agostino & Russell, 2005), as well as by visual inspection of population separation on the PCs. The PCs that reflected population structure (PC1 for Coho and Sockeye; PCs 1-4 for Chinook and Steelhead; Figures S1-S18), as well as their interactions with the SNP, were then included as covariates in the models.

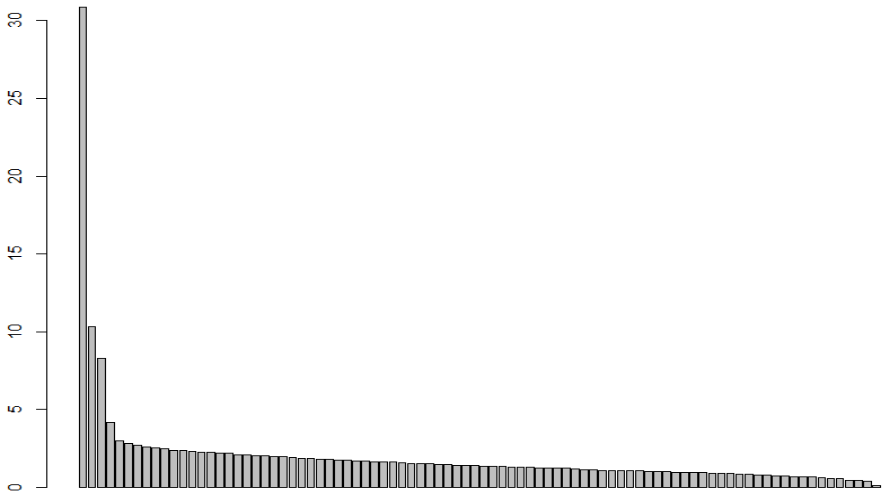


Figure S1. Plot of eigenvalues from the principal component analysis conducted using 86 putatively neutral loci for Chinook Salmon (four loci were removed due to high rate of missing genotypes). The first four PCs reflected population structure, as determined by the scree test and through visual inspection of population separation along the PCs.

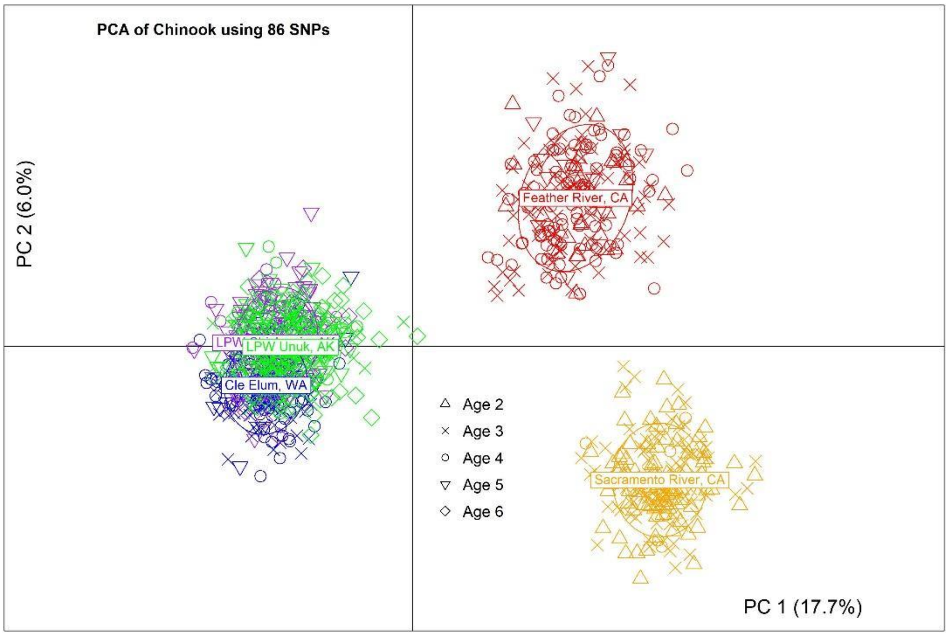


Figure S2. Plot of Chinook Salmon by population and age along PC1 and PC2 from the principal component analysis conducted using 86 putatively neutral loci.

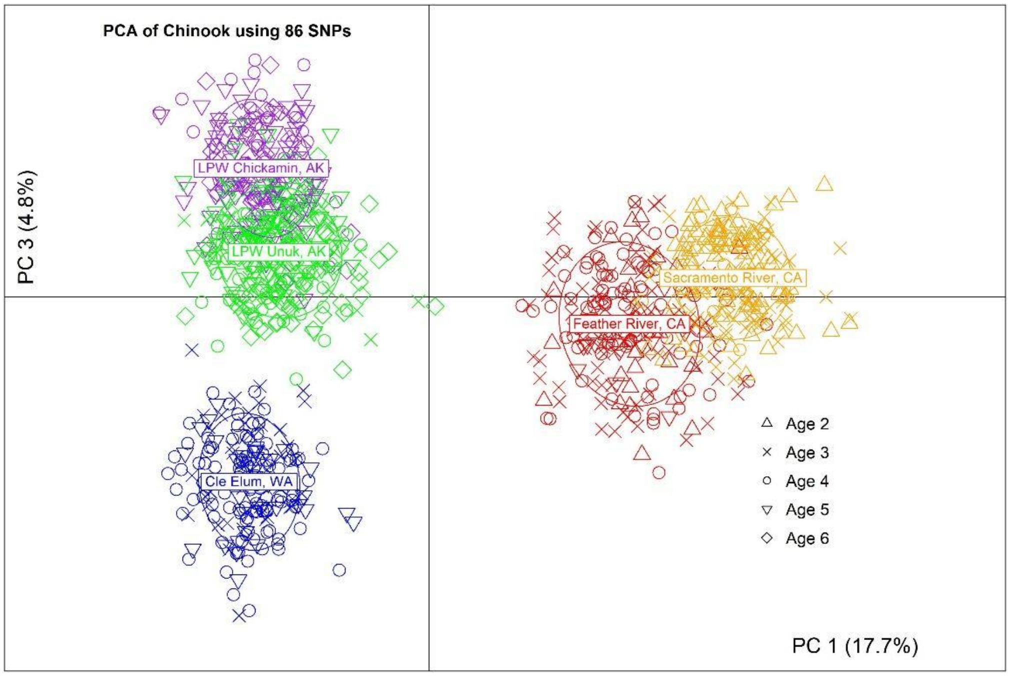


Figure S3. Plot of Chinook Salmon by population and age along PC1 and PC3 from the principal component analysis conducted using 86 putatively neutral loci.

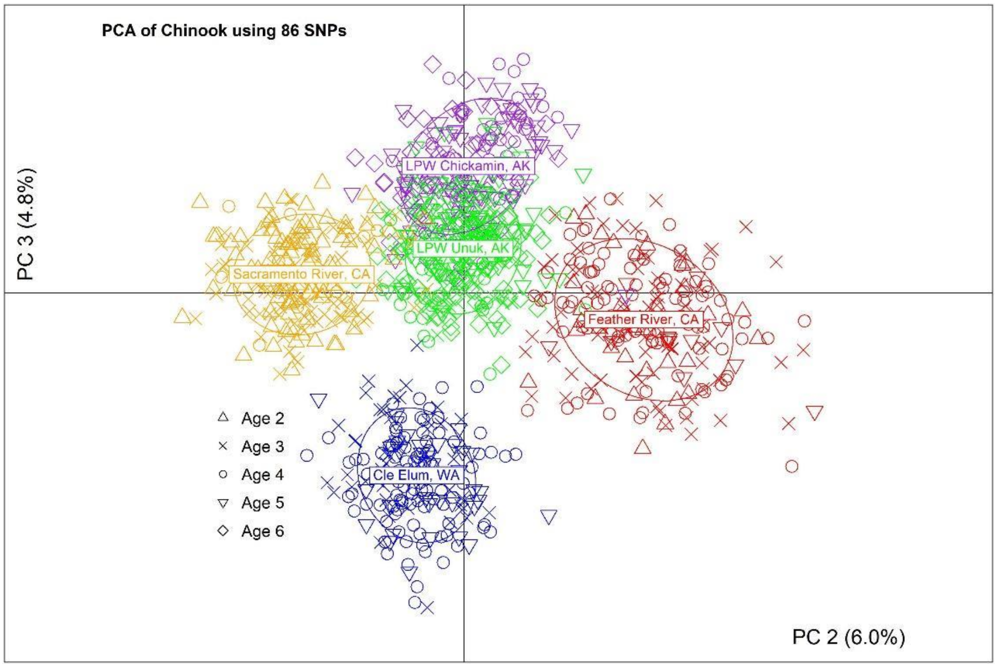


Figure S4. Plot of Chinook Salmon by population and age along PC2 and PC3 from the principal component analysis conducted using 86 putatively neutral loci.

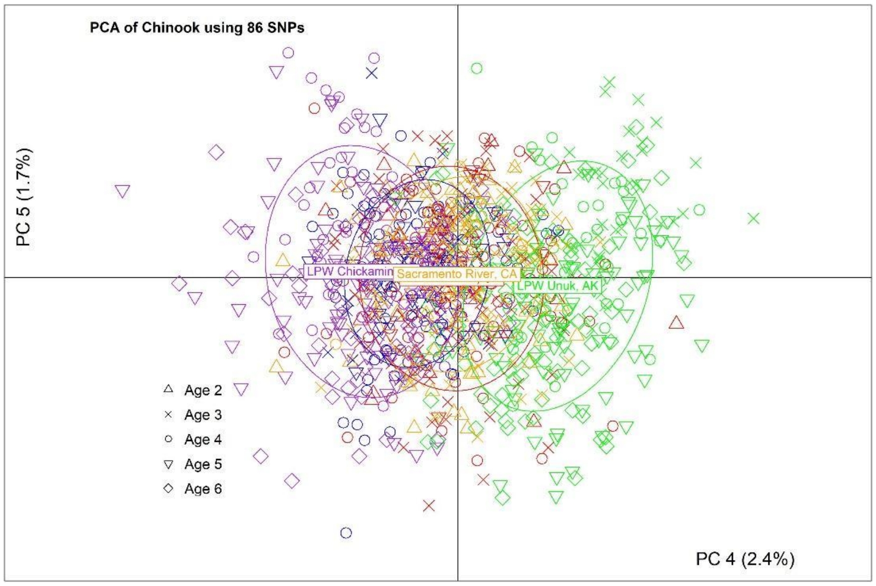


Figure S5. Plot of Chinook Salmon by population and age along PC4 and PC5 from the principal component analysis conducted using 86 putatively neutral loci.

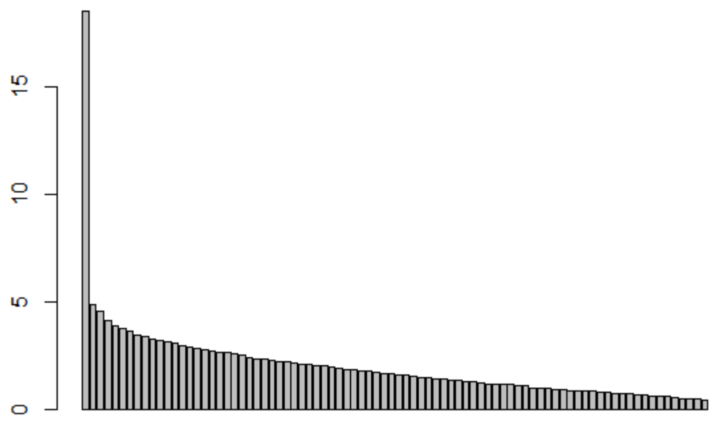


Figure S6. Plot of eigenvalues from the principal component analysis conducted using 85 putatively neutral loci for Coho Salmon (84 polymorphic loci and 1 monomorphic locus). The first PC reflected population structure, as determined by the scree test and through visual inspection of population separation along the PCs.

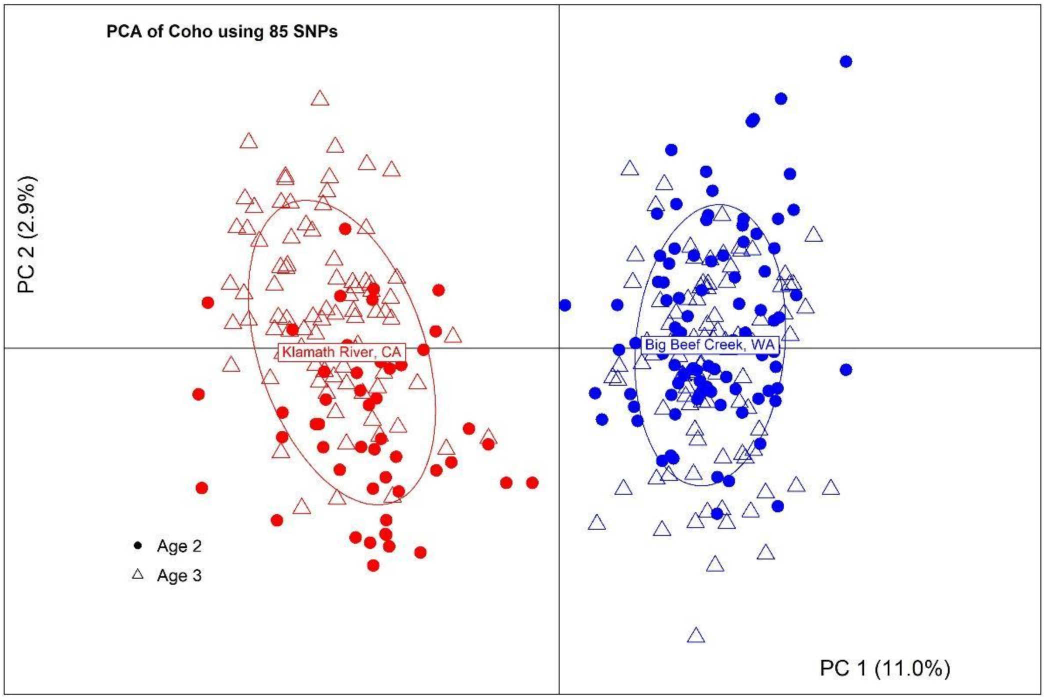


Figure S7. Plot of Coho Salmon by population and age along PC1 and PC2 from the principal component analysis conducted using 85 putatively neutral loci.

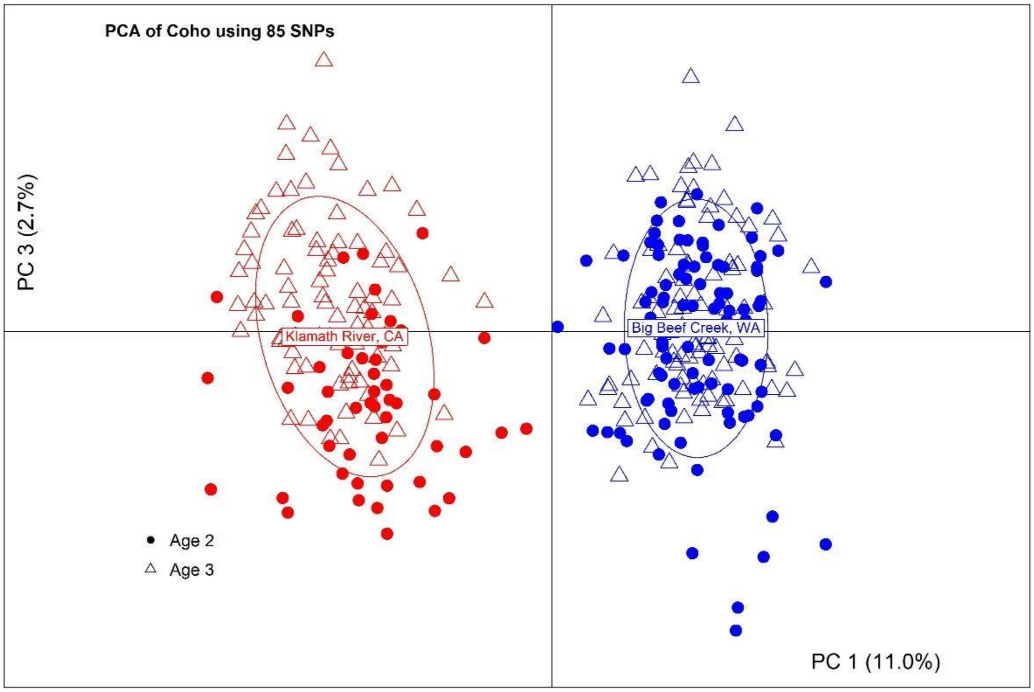


Figure S8. Plot of Coho Salmon by population and age along PC1 and PC3 from the principal component analysis conducted using 85 putatively neutral loci.

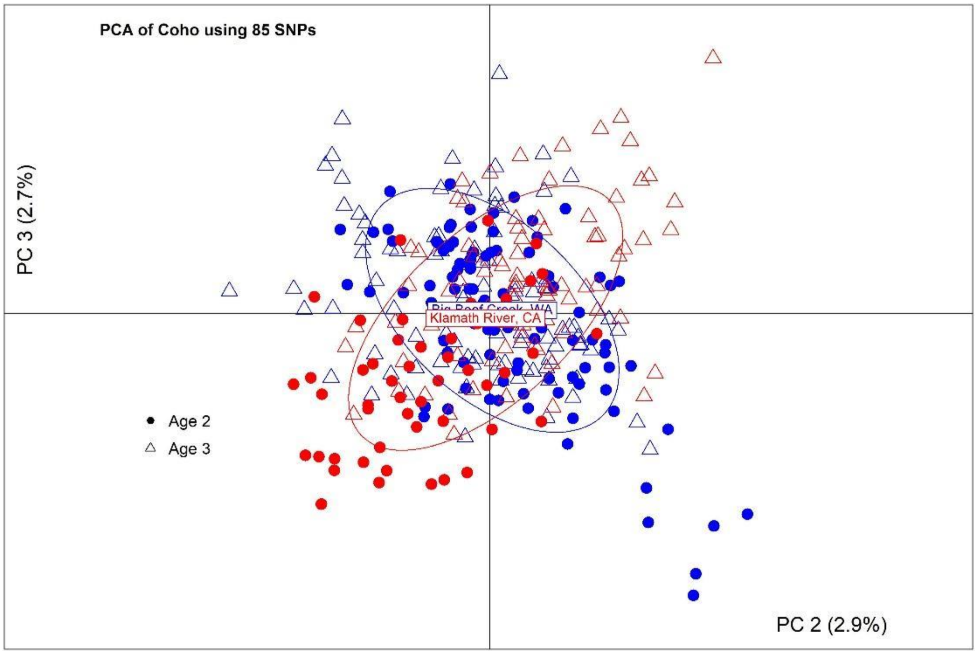


Figure S9. Plot of Coho Salmon by population and age along PC2 and PC3 from the principal component analysis conducted using 85 putatively neutral loci.

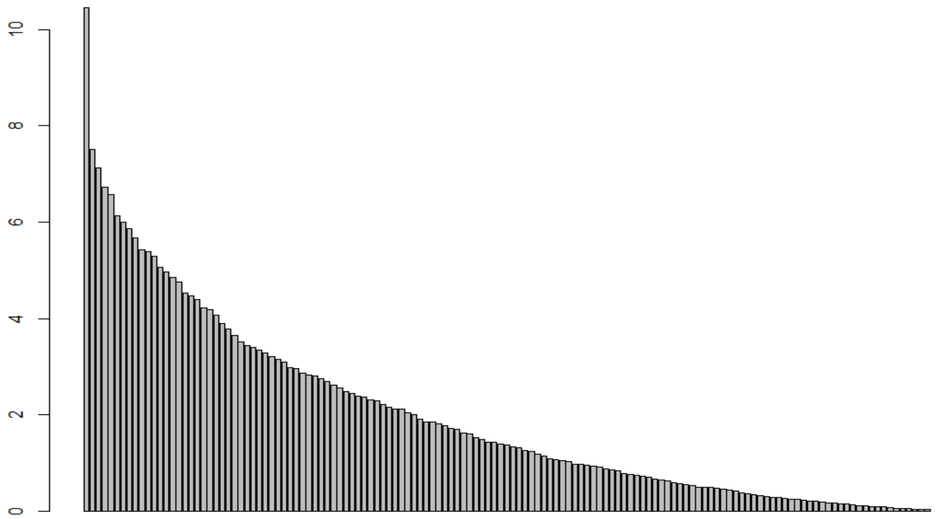


Figure S10. Plot of eigenvalues from the principal component analysis conducted using 137 of the putatively neutral loci for Sockeye Salmon (other loci excluded from PCA as they may not be completely neutral). The first PC reflected population structure, as determined by the scree test and through visual inspection of population separation along the PCs.

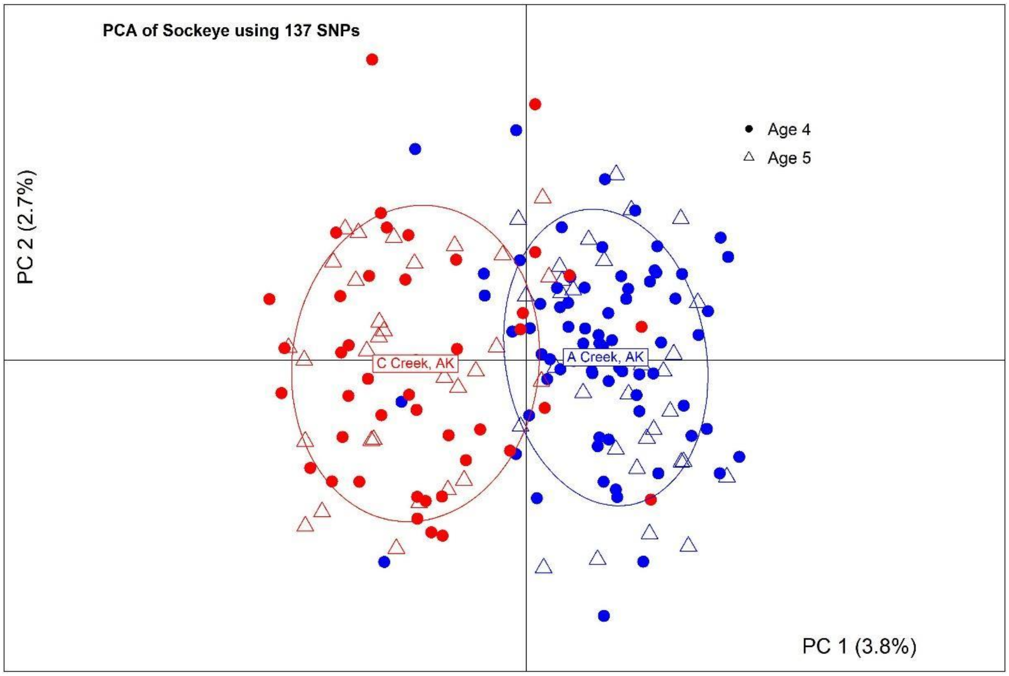


Figure S11. Plot of Sockeye Salmon by population and age along PC1 and PC2 from the principal component analysis conducted using 137 putatively neutral loci.

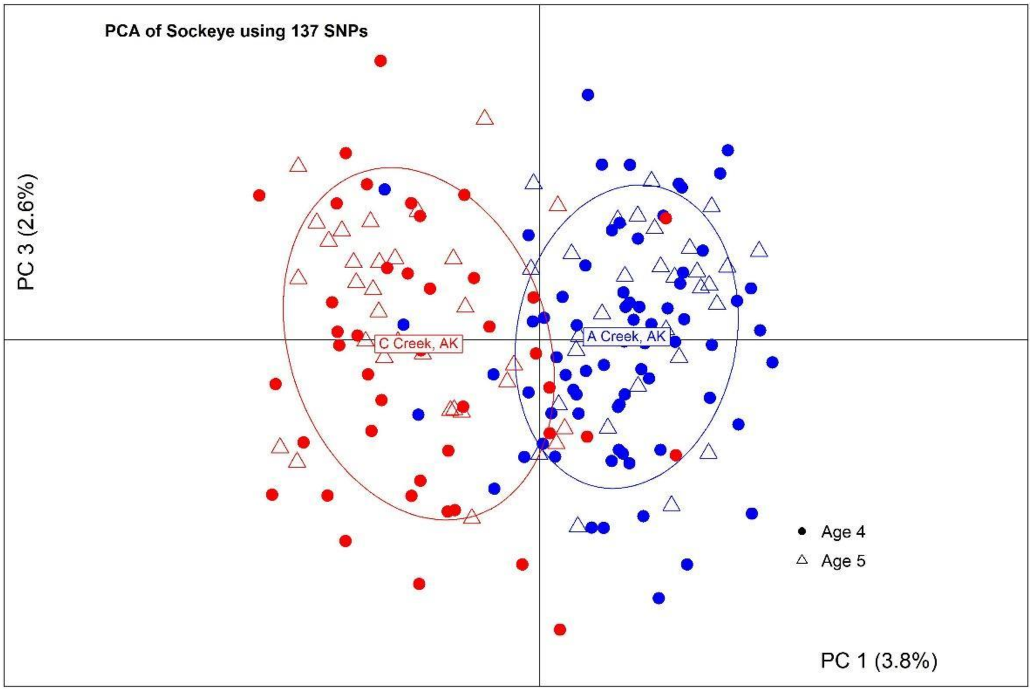


Figure S12. Plot of Sockeye Salmon by population and age along PC1 and PC3 from the principal component analysis conducted using 137 putatively neutral loci.

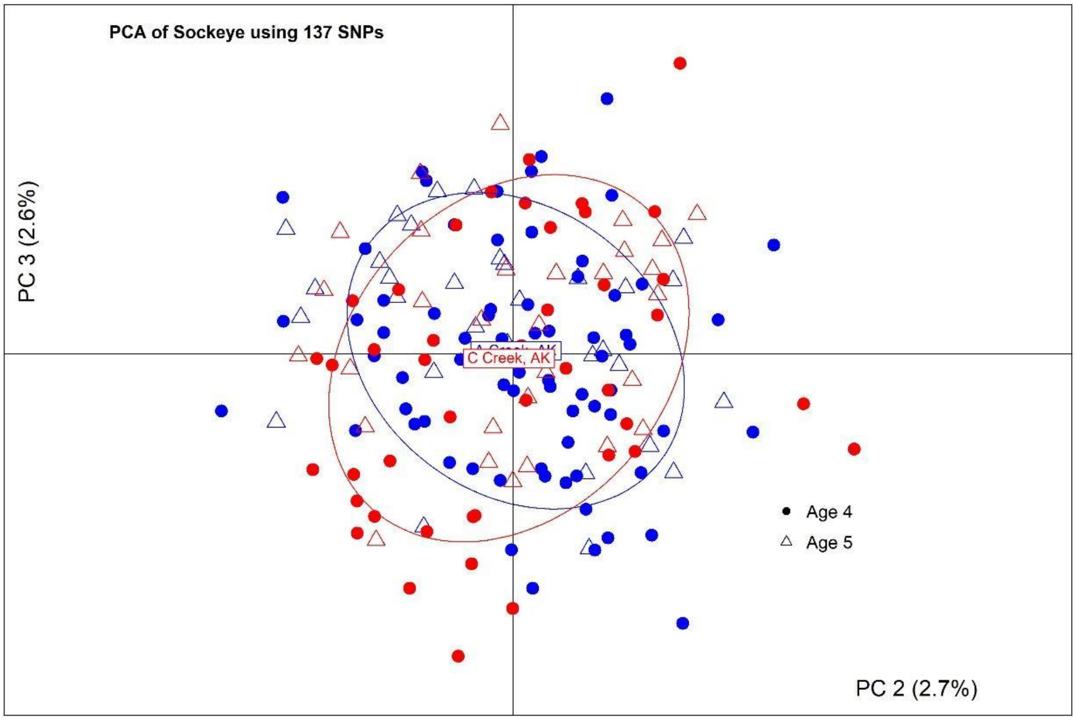


Figure S13. Plot of Sockeye Salmon by population and age along PC2 and PC3 from the principal component analysis conducted using 137 putatively neutral loci.

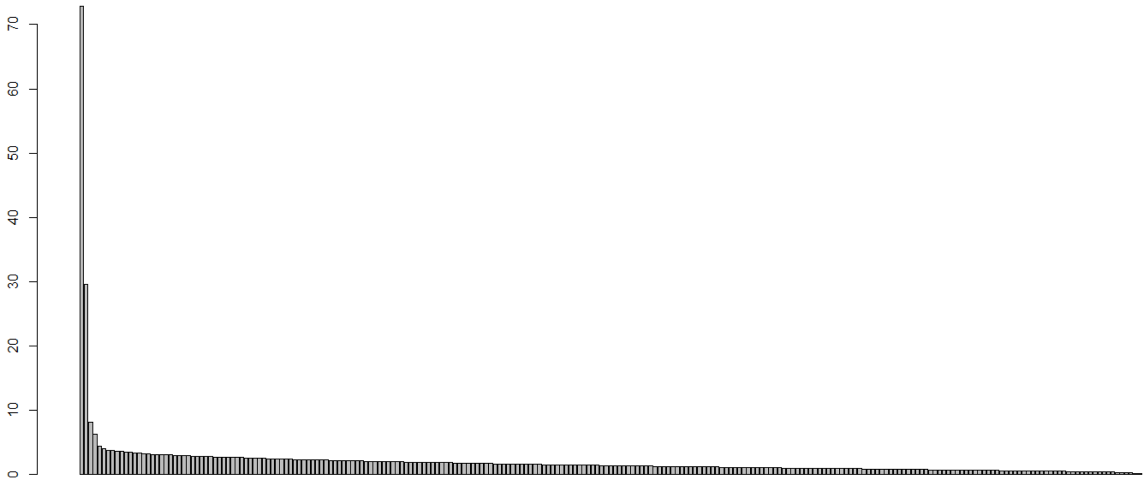


Figure S14. Plot of eigenvalues from the principal component analysis conducted using 239 of the putatively neutral loci for Steelhead Trout (three loci excluded). The first four PCs reflected population structure, as determined by the scree test and through visual inspection of population separation along the PCs.

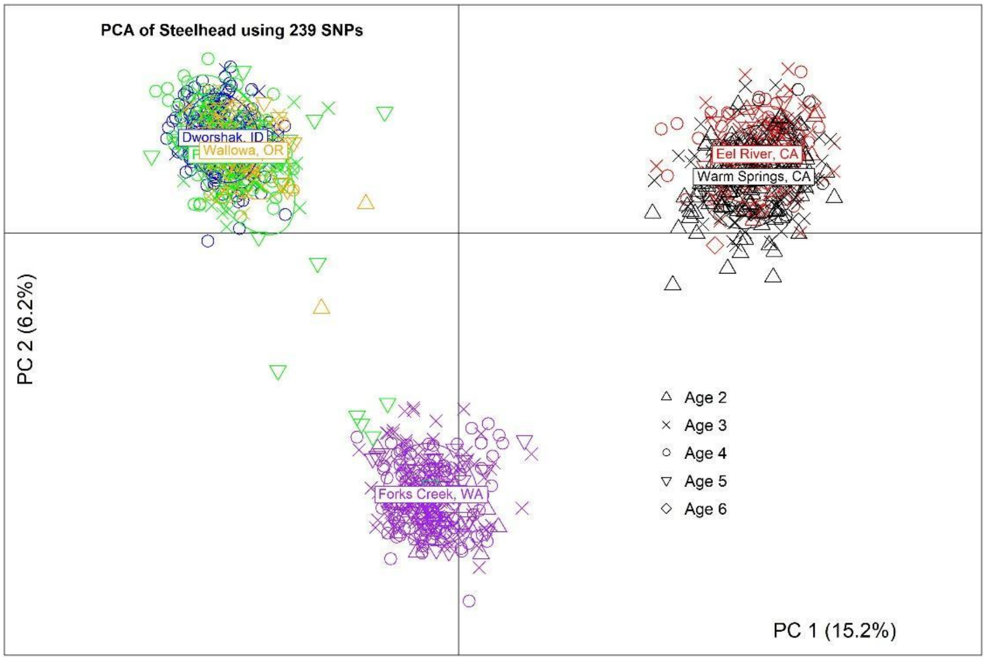


Figure S15. Plot of Steelhead Trout by population and age along PC1 and PC2 from the principal component analysis conducted using 239 putatively neutral loci.

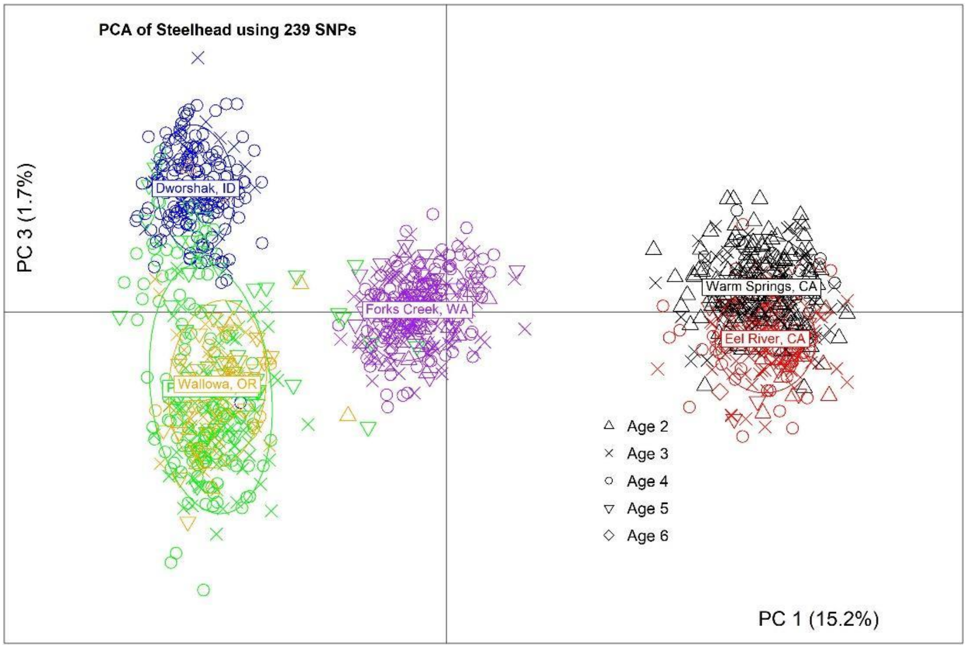


Figure S16. Plot of Steelhead Trout by population and age along PC1 and PC3 from the principal component analysis conducted using 239 putatively neutral loci.

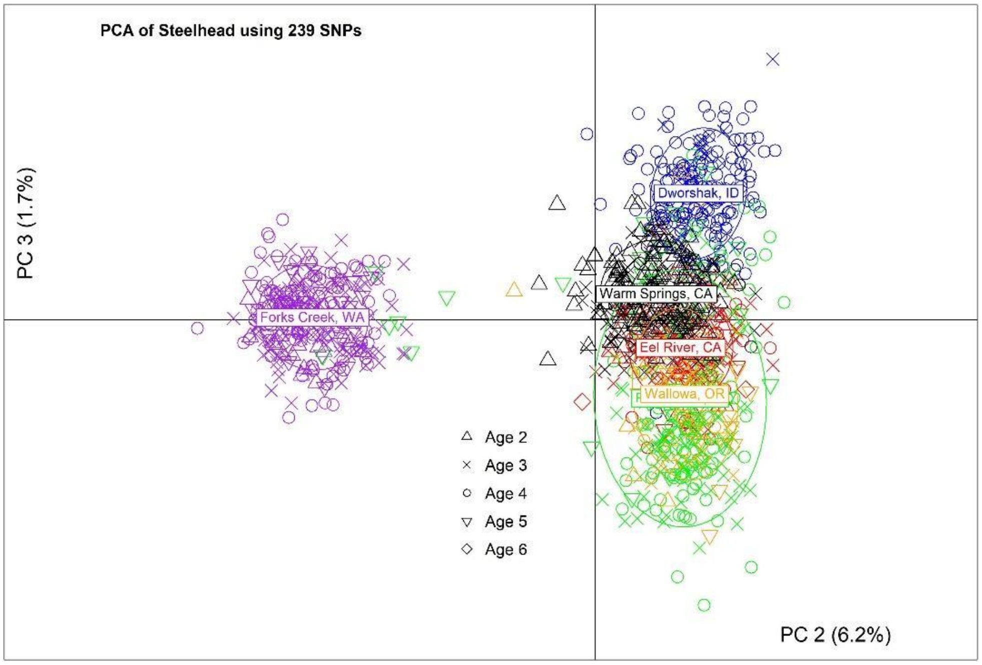


Figure S17. Plot of Steelhead Trout by population and age along PC2 and PC3 from the principal component analysis conducted using 239 putatively neutral loci.

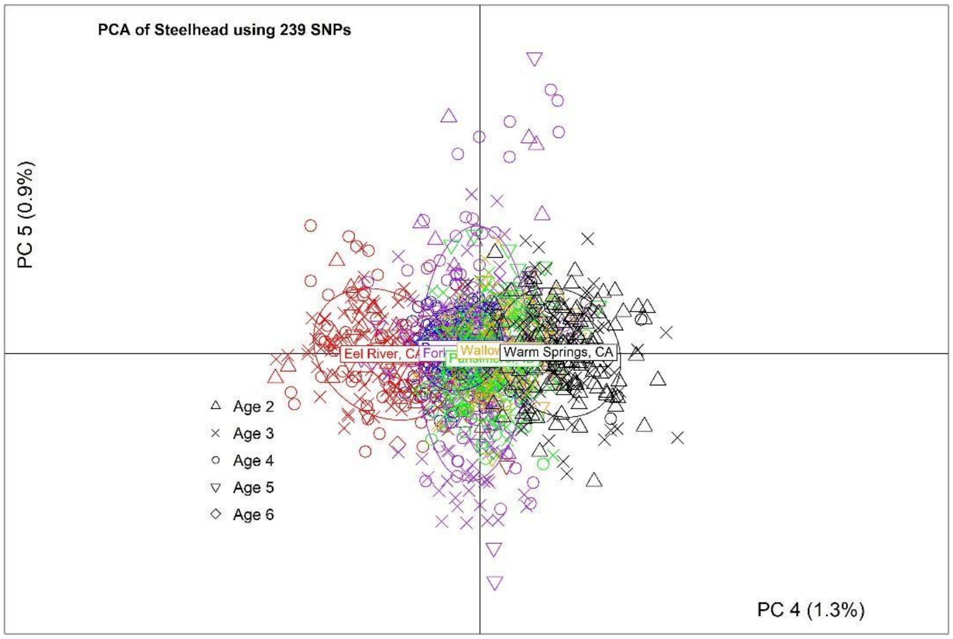


Figure S18. Plot of Steelhead Trout by population and age along PC4 and PC5 from the principal component analysis conducted using 239 putatively neutral loci.

*Pairwise relatedness*

To determine whether relatedness might have influenced the results, molecular estimates of pairwise relatedness between samples within each population was estimated using putatively neutral loci and the R package *related* (v. 0.8, Pew, Muir, Wang, & Frasier, 2015), the R implementation of the COANCESTRY program (Wang, 2011). The program implements seven relatedness estimators and enables users to perform simulations to determine the best estimator for their data. For each population, parent-offspring, full sibling, half sibling, and unrelated relationships (n=100 per category) were simulated; simulations were parameterized by the number of loci and the allele frequencies of those loci from the empirical data sets. Inbreeding was accounted for in the two likelihood estimators of relatedness, and 100 reference individuals were used when estimating relatedness with the triadic likelihood method (Milligan, 2003; Wang, 2007). The estimator that had the highest correlation between the true and estimated values from each simulation was then used to obtain empirical estimates of pairwise relatedness for each population. The DyadML and TrioML methods had the highest correlations between true and estimated values for all populations (all >0.89). These molecular estimates confirmed that samples within populations comprised mostly unrelated individuals (Figures S19-S33).

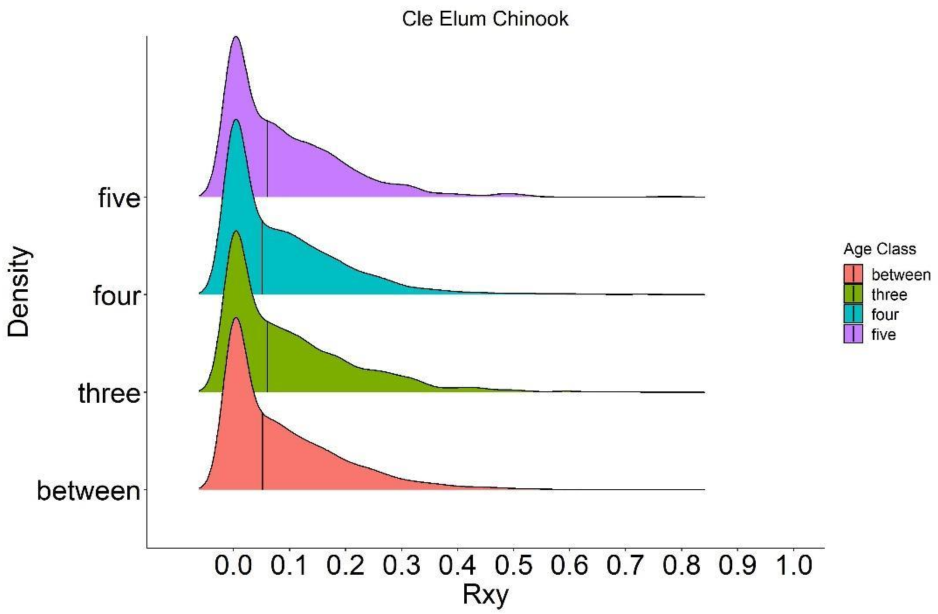


Figure S19. Density plot of estimates of pairwise relatedness, $R_{xy}$, within and between age classes of Cle Elum Chinook Salmon obtained from 86 putatively neutral loci (four loci were removed due to high rate of missing genotypes) using the DyadML method. Vertical black lines denote the median of the respective distributions.

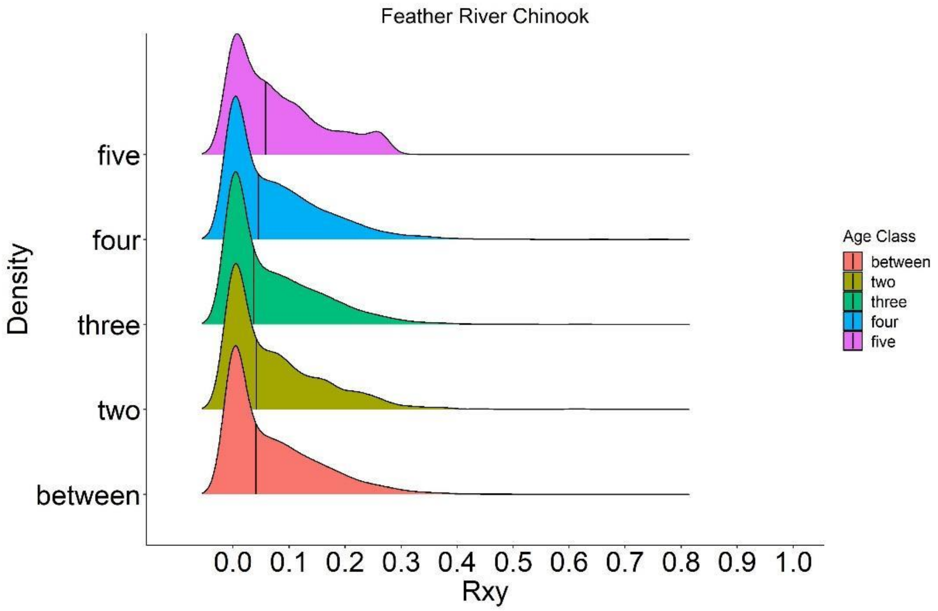


Figure S20. Density plot of estimates of pairwise relatedness, $R_{xy}$, within and between age classes of Feather River Chinook Salmon obtained from 86 putatively neutral loci using the TrioML method. Vertical black lines denote the median of the respective distributions.

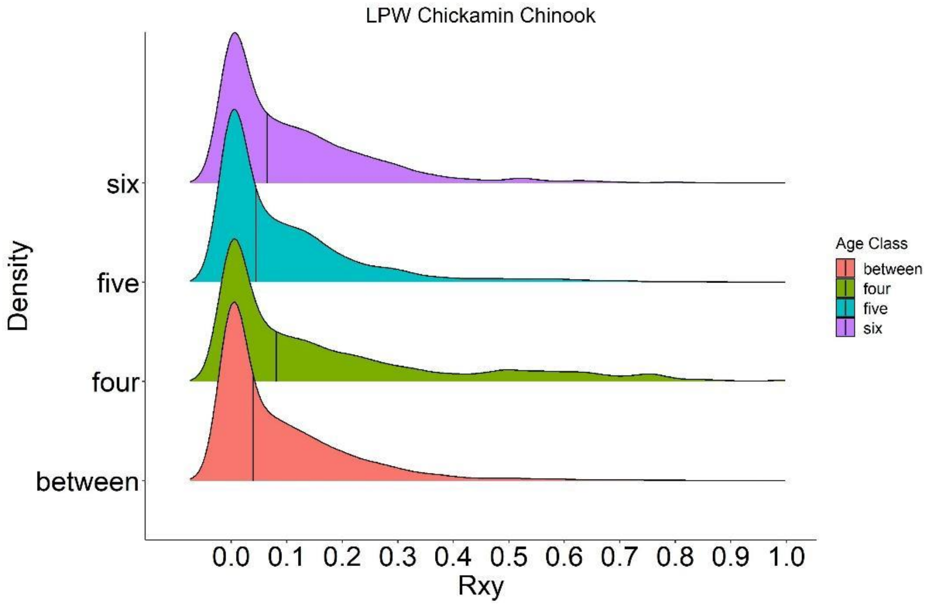


Figure S21. Density plot of estimates of pairwise relatedness, $R_{xy}$, within and between age classes of Little Port Walter Chickamin Stock Chinook Salmon obtained from 86 putatively neutral loci using the TrioML method. Vertical black lines denote the median of the respective distributions.

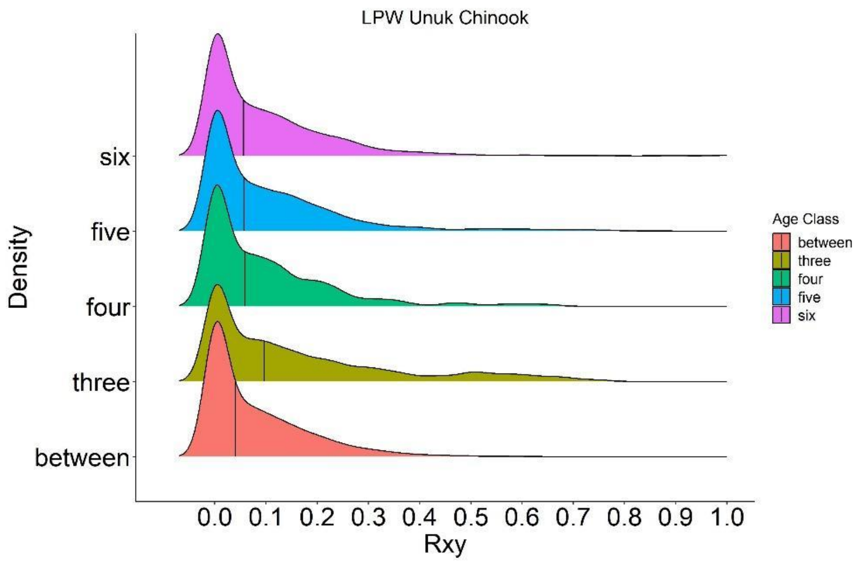


Figure S22. Density plot of estimates of pairwise relatedness, $R_{xy}$, within and between age classes of Little Port Walter Unuk Stock Chinook Salmon obtained from 86 putatively neutral loci using the TrioML method. Vertical black lines denote the median of the respective distributions.

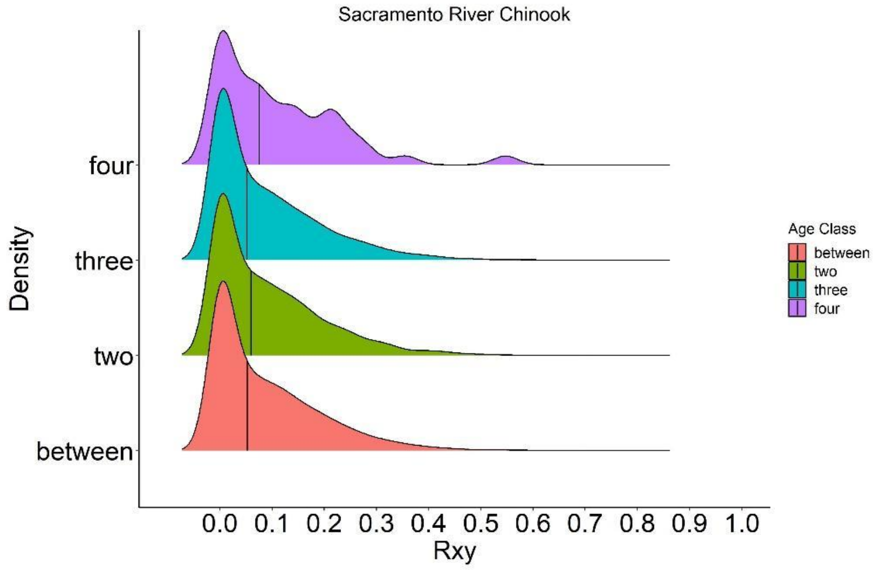


Figure S23. Density plot of estimates of pairwise relatedness, $R_{xy}$, within and between age classes of Sacramento River Chinook Salmon obtained from 86 putatively neutral loci using the DyadML method. Vertical black lines denote the median of the respective distributions.

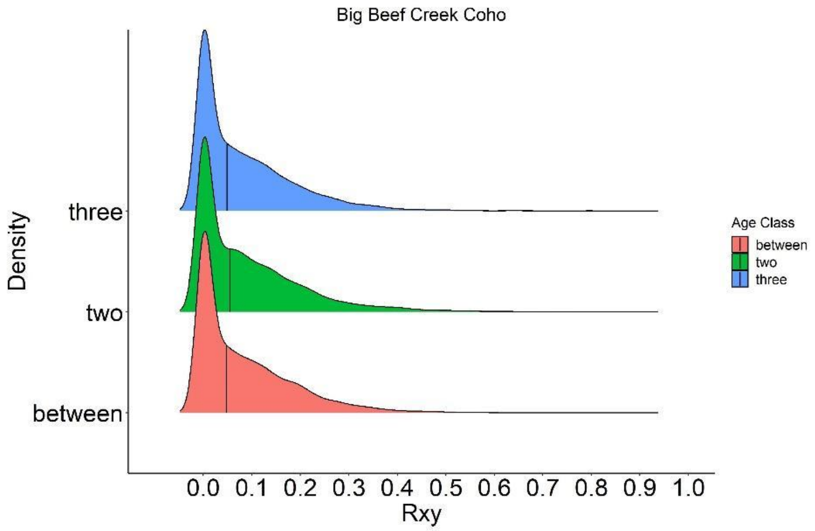


Figure S24. Density plot of estimates of pairwise relatedness, $R_{xy}$, within and between age classes of Big Beef Creek Coho Salmon obtained from 85 putatively neutral loci (84 polymorphic loci and 1 monomorphic locus) using the DyadML method. Vertical black lines denote the median of the respective distributions.

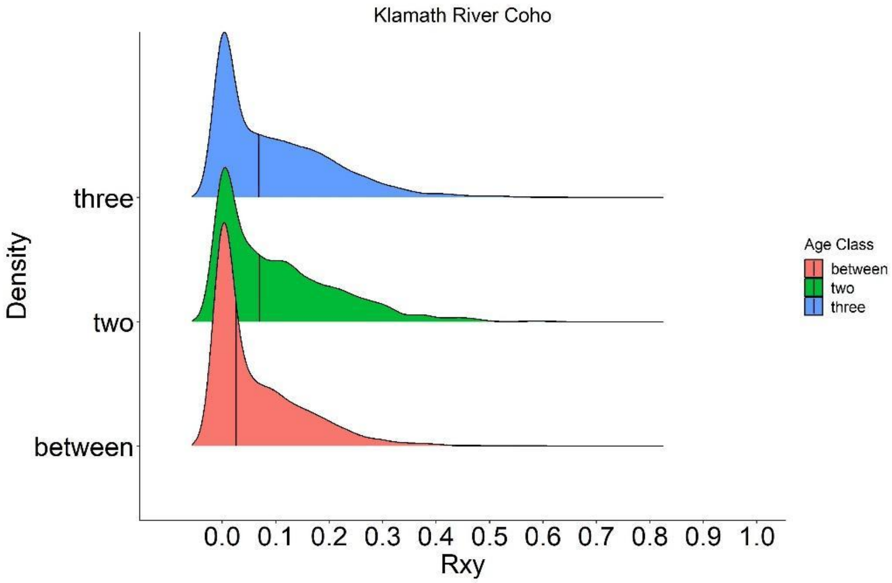


Figure S25. Density plot of estimates of pairwise relatedness, $R_{xy}$, within and between age classes of Klamath River Coho Salmon obtained from 85 putatively neutral loci using the TrioML method. Vertical black lines denote the median of the respective distributions.

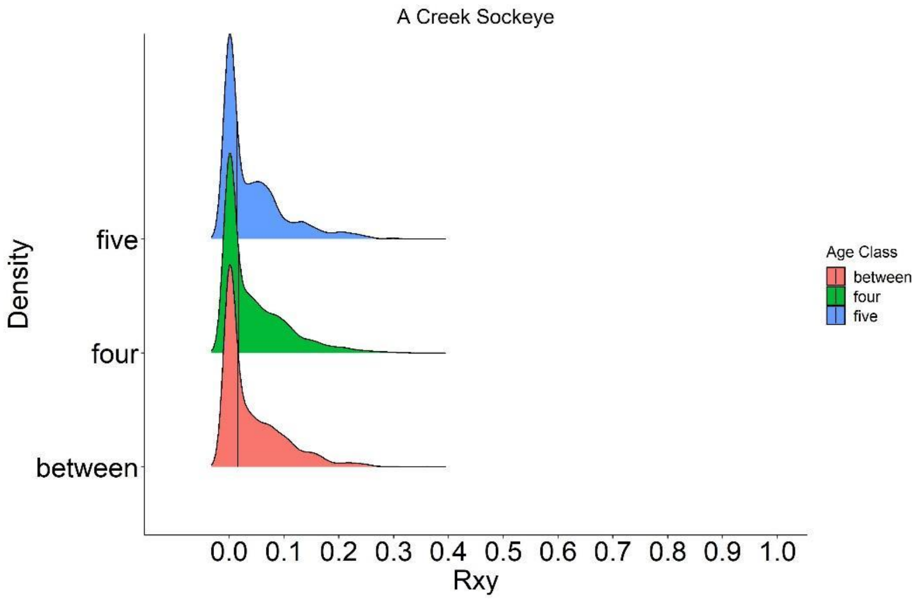


Figure S26. Density plot of estimates of pairwise relatedness, $R_{xy}$, within and between age classes of A Creek Sockeye Salmon obtained from 137 putatively neutral loci (other loci excluded as they may not be completely neutral) using the DyadML method. Vertical black lines denote the median of the respective distributions.

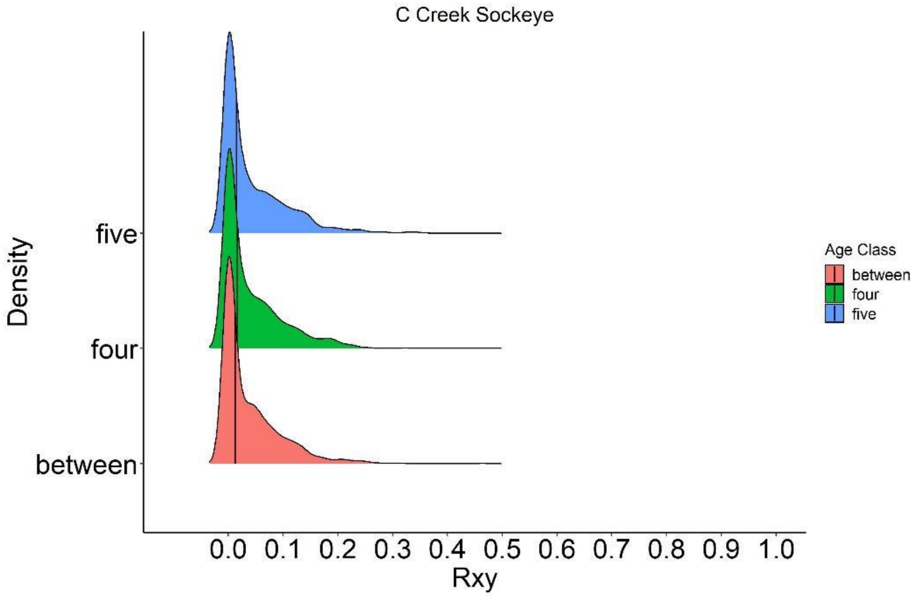


Figure S27. Density plot of estimates of pairwise relatedness, $R_{xy}$, within and between age classes of C Creek Sockeye Salmon obtained from 137 putatively neutral loci using the TrioML method. Vertical black lines denote the median of the respective distributions.

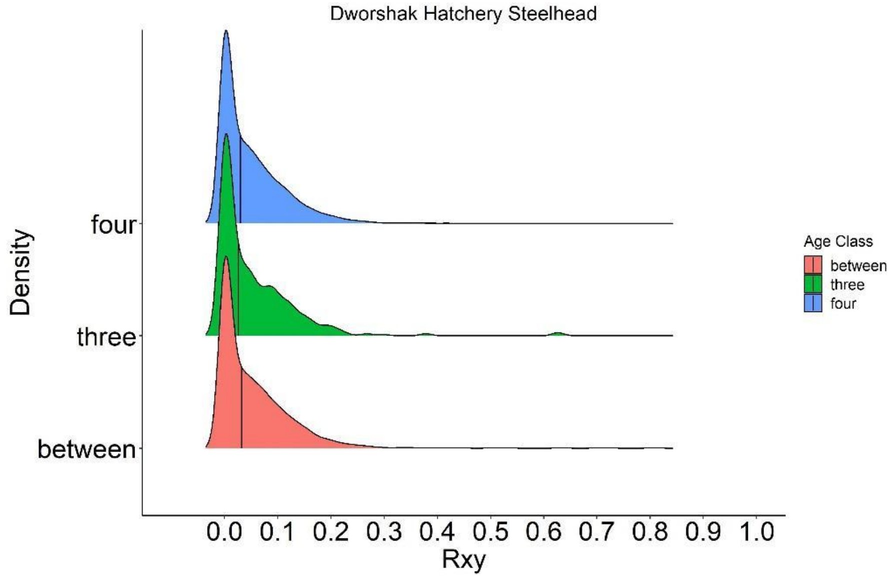


Figure S28. Density plot of estimates of pairwise relatedness, $R_{xy}$, within and between age classes of Dworshak Hatchery Steelhead Trout obtained from 239 putatively neutral loci (three loci excluded) using the DyadML method. Vertical black lines denote the median of the respective distributions.

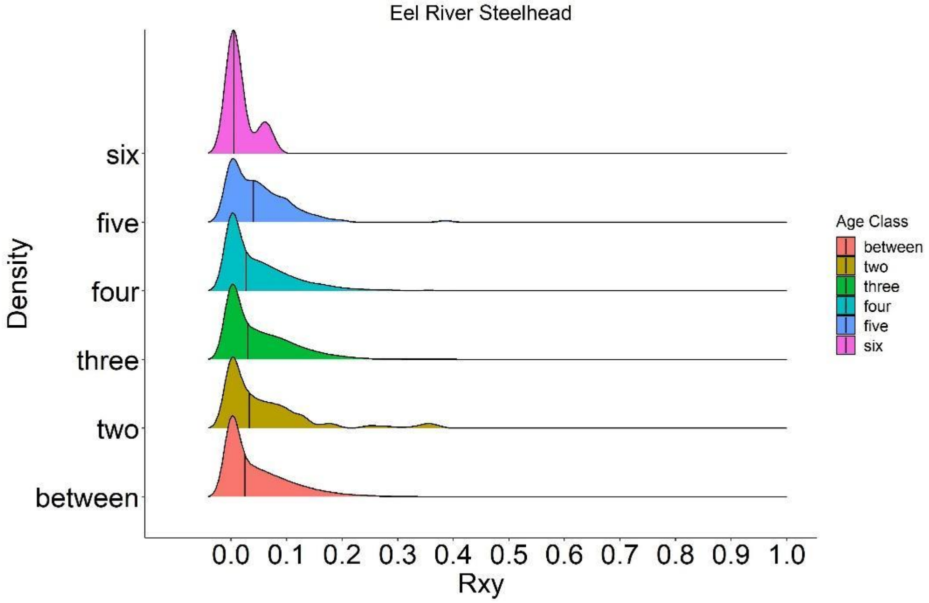


Figure S29. Density plot of estimates of pairwise relatedness, $R_{xy}$, within and between age classes of Eel River Steelhead Trout obtained from 239 putatively neutral loci using the DyadML method. Vertical black lines denote the median of the respective distributions.

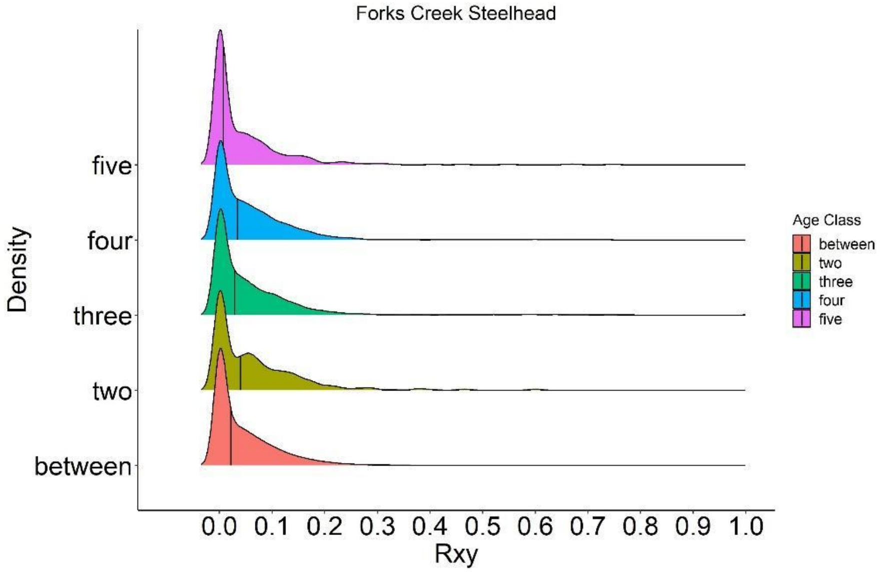


Figure S30. Density plot of estimates of pairwise relatedness, $R_{xy}$, within and between age classes of Forks Creek Steelhead Trout obtained from 239 putatively neutral loci using the DyadML method. Vertical black lines denote the median of the respective distributions.

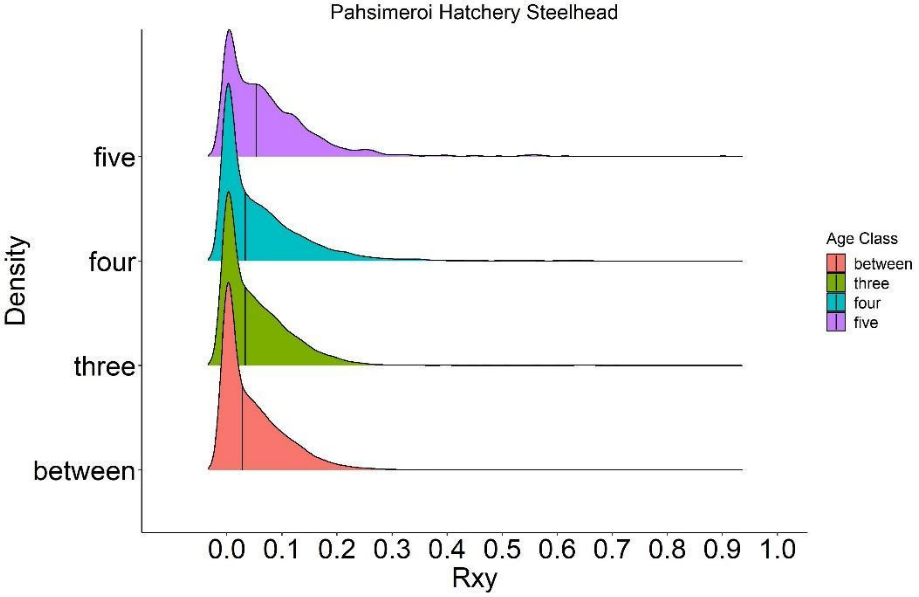


Figure S31. Density plot of estimates of pairwise relatedness, $R_{xy}$, within and between age classes of Pahsimeroi Hatchery Steelhead Trout obtained from 239 putatively neutral loci using the DyadML method. Vertical black lines denote the median of the respective distributions.

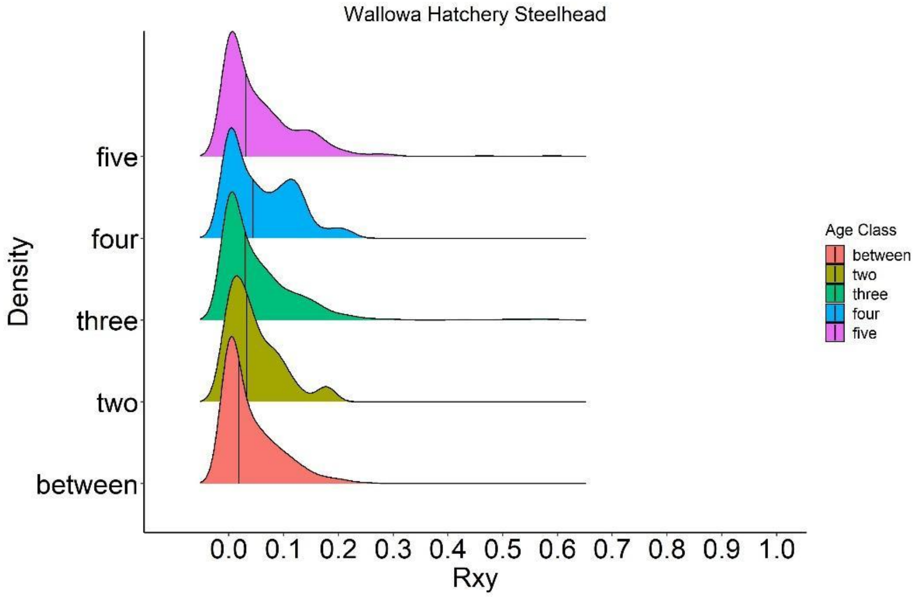


Figure S32. Density plot of estimates of pairwise relatedness, $R_{xy}$, within and between age classes of Wallowa Hatchery Steelhead Trout obtained from 239 putatively neutral loci using the TrioML method. Vertical black lines denote the median of the respective distributions.

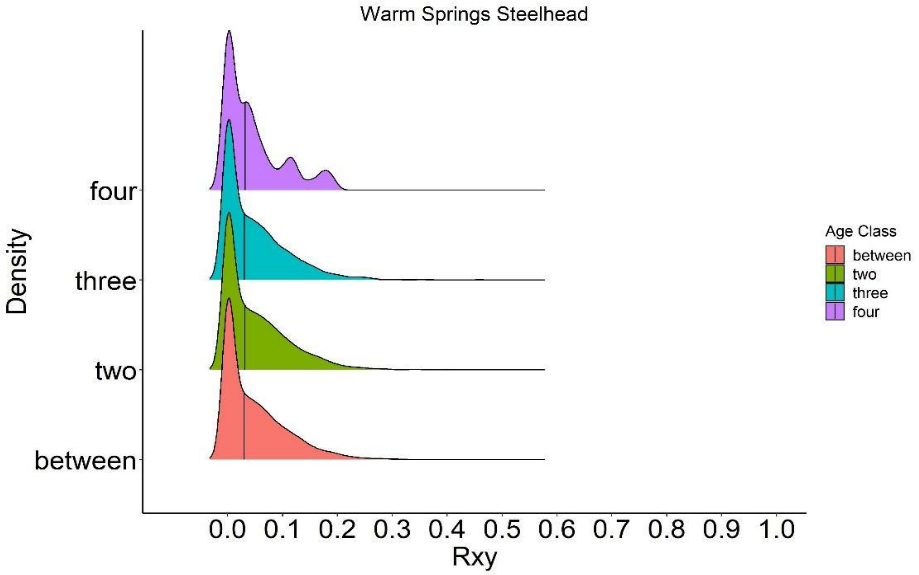


Figure S33. Density plot of estimates of pairwise relatedness, $R_{xy}$, within and between age classes of Warm Springs Hatchery Steelhead Trout obtained from 239 putatively neutral loci using the DyadML method. Vertical black lines denote the median of the respective distributions.

### Literature Cited


Abadía-Cardoso, A., Anderson, E. C., Pearse, D. E., & Garza, J. C. (2013). Large-scale parentage analysis reveals reproductive patterns and heritability of spawn timing in a hatchery population of steelhead (*Oncorhynchus mykiss*). *Molecular Ecology, 22*(18), 4733-4746. doi:10.1111/mec.12426

Bowersox, B. J., Corsi, M. P., McCormick, J. L., Copeland, T., & Campbell, M. R. (2019). Examining life history shifts and genetic composition in a hatchery steelhead population, with implications for fishery and ocean selection. *Transactions of the American Fisheries Society, 148*(6), 1056-1068. doi:10.1002/tafs.10199

Cattell, R. B. (1966). The scree test for the number of factors. *Multivariate Behavioral Research, 1*(2), 245-276.

Clemento, A. (2013). *Creation and utilization of novel genetic methods for studying and improving management of Chinook salmon populations.* (PhD Dissertation), University of California Santa Cruz, Santa Cruz, California.

Clutter, R. I., & Whitesel, L. E. (1956). *Collection and interpretation of Sockeye Salmon scales*. International Pacific Salmon Fisheries Commission.

Copeland, T., Ackerman, M. W., Wright, K. K., & Byrne, A. (2017). Life history diversity of Snake River steelhead populations between and within management categories. *North American Journal of Fisheries Management, 37*(2), 395-404. doi:10.1080/02755947.2016.1264506

Crawford, B. A. (1979). The origin and history of the trout brood stocks of the Washington Department of Game. Washington State Game Department, Fishery Research Report SH167.T86C7. Olympia.

D'agostino, R. B., & Russell, H. K. (2005). Scree test. In *Encyclopedia of Biostatistics*.

Hess, J. E., Ackerman, M. W., Fryer, J. K., Hasselman, D. J., Steele, C. A., Stephenson, J. J., . . . Narum, S. R. (2016). Differential adult migration-timing and stock-specific abundance of steelhead in mixed stock assemblages. *Ices Journal of Marine Science, 73*(10), 2606-2615. doi:10.1093/icesjms/fsw138

Jombart, T. (2008). *adegenet:* a R package for the multivariate analysis of genetic markers. *Bioinformatics, 24*(11), 1403-1405. doi:10.1093/bioinformatics/btn129

Jombart, T., & Ahmed, I. (2011). adegenet 1.3-1: new tools for the analysis of genome-wide SNP data. *Bioinformatics, 27*(21), 3070-3071. doi:10.1093/bioinformatics/btr521

Knudsen, C. M., Schroder, S. L., Busack, C. A., Johnston, M. V., Pearsons, T. N., Bosch, W. J., & Fast, D. E. (2006). Comparison of life history traits between first-generation hatchery and wild upper Yakima river spring Chinook salmon. *Transactions of the American Fisheries Society, 135*(4), 1130-1144. doi:10.1577/t05-121.1

Kodama, M., Hard, J. J., & Naish, K. A. (2012). Temporal variation in selection on body length and date of return in a wild population of coho salmon, *Oncorhynchus kisutch*. *Bmc Evolutionary Biology, 12*, 12. doi:10.1186/1471-2148-12-116

Lin, J., Quinn, T. P., Hilborn, R., & Hauser, L. (2008). Fine-scale differentiation between sockeye salmon ecotypes and the effect of phenotype on straying. *Heredity, 101*(4), 341-350. doi:10.1038/hdy.2008.59

May, S. A., McKinney, G. J., Hilborn, R., Hauser, L., & Naish, K. (2020). Power of a dual-use SNP panel for pedigree reconstruction and population assignment. *Ecology and Evolution*(Accepted).

Milligan, B. G. (2003). Maximum-likelihood estimation of relatedness. *Genetics, 163*(3), 1153-1167.

Myers, J. M., Kope, R. G., Bryant, G. J., Teel, D., Lierheimer, L. J., Wainwright, T. C., . . . Waples, R. S. (1998). *Status review of Chinook salmon from Washington, Idaho, Oregon, and California*. NOAA Tech. Memo.

Naish, K. A., Seamons, T. R., Dauer, M. B., Hauser, L., & Quinn, T. P. (2013). Relationship between effective population size, inbreeding and adult fitness-related traits in a steelhead (*Oncorhynchus mykiss*) population released in the wild. *Molecular Ecology, 22*(5), 1295-1309. doi:10.1111/mec.12185

Peterson, D. A., Hilborn, R., & Hauser, L. (2014). Local adaptation limits lifetime reproductive success of dispersers in a wild salmon metapopulation. *Nature Communications, 5*, 7. doi:10.1038/ncomms4696

Pew, J., Muir, P. H., Wang, J. L., & Frasier, T. R. (2015). related: an R package for analysing pairwise relatedness from codominant molecular markers. *Molecular Ecology Resources, 15*(3), 557-561. doi:10.1111/1755-0998.12323

Starks, H. (2014). *Genetic pedigree inference in coho salmon: A powerful tool for guiding the management of an ESA-listed species.* (MS Thesis), University of California Santa Cruz, Santa Cruz, California.

Templin, W. D. (2001). The history of propagation and transportation of Chinook salmon *Oncorhynchus tshawytscha* stocks at hatcheries in Southeast Alaska, 1972-1998. Regional Information Report No. 5J01-05. Alaska Department of Fish and Game.

Wang, J. (2007). Triadic IBD coefficients and applications to estimating pairwise relatedness. *Genetics Research, 89*(3), 135-153. doi:10.1017/s0016672307008798

Wang, J. L. (2011). COANCESTRY: a program for simulating, estimating and analysing relatedness and inbreeding coefficients. *Molecular Ecology Resources, 11*(1), 141-145. doi:10.1111/j.1755-0998.2010.02885.x

Waters, C. D., Hard, J. J., Brieuc, M. S. O., Fast, D. E., Warheit, K. I., Knudsen, C. M., . . . Naish, K. A. (2018). Genomewide association analyses of fitness traits in captive-reared Chinook salmon: Applications in evaluating conservation strategies. *Evolutionary Applications, 00*, 1-16. doi:10.1111/eva.12599

Waters, C. D., Hard, J. J., Brieuc, M. S. O., Fast, D. E., Warheit, K. I., Waples, R. S., . . . Naish, K. A. (2015). Effectiveness of managed gene flow in reducing genetic divergence associated with captive breeding. *Evolutionary Applications, 8*(10), 956-971. doi:10.1111/eva.12331
